# Hepatocyte ERα orchestrates sex-specific liver adaptation to fasting and feeding states

**DOI:** 10.64898/2025.12.05.692559

**Authors:** Blandine Tramunt, Marie-Lou Calmy, Arnaud Polizzi, Marine Huillet, Naia Grandgeorge, Ana Avalrez-Tena, Sarra Smati, Tiffany Fougeray, Frédéric Boudou, Marion Régnier, Anne Fougerat, Stéphanie Gayral, Frédéric Lasserre, Claire Naylies, Yannick Lippi, Christophe Bureau, Maéva Guillaume, Dominique Langin, Atish Mukherji, Raphael Métivier, Yuna Blum, Nicolas Loiseau, Walter Wahli, Hélène Duez, Catherine Postic, Hervé Guillou, Pierre Gourdy, Alexandra Montagner

**Affiliations:** Toulouse University, Institute of Metabolic and Cardiovascular Diseases (I2MC), INSERM, Toulouse, France; Toulouse University, CHU de Toulouse, Department of Diabetology, Metabolic Diseases and Nutrition, Toulouse, France; Toulouse University, Research Center in Food toxicology (ToxAlim), INRAe ENVT, INP-Purpan, UMR 1331, Toulouse University, Toulouse, France; Paris Cité University, Institut Cochin, CNRS, INSERM, Paris, France; Toulouse University, CHU de Toulouse, Department of Gastroenterology and Hepatology, Toulouse, France; Clinique Pasteur, Department of Gastroenterology and Hepatology, Toulouse; Strasbourg University, Institute for Translational Medecine and Liver Disease, Strasbourg, France; Institute Genetics and Development of Rennes (IGRD), UMR 6290 CNRS/UR – ERL U1305 INSERM, Rennes University, Rennes, France; Center for Integrative Genomics (CIG), Lausanne University, Le Génopode, Lausanne, Switzerland; Lee Kong Chian School of Medicine, Nanyang Technological University Singapore, Singapore, Singapore; Lille University, INSERM, CHU Lille, Institut Pasteur of Lille, U1011-EGID, France

## Abstract

Sex differences in hepatic physiology are thought to influence individual susceptibility to chronic liver diseases including Metabolic-Associated Steatotic Liver Disease (MALSD). Previous studies demonstrated the central contribution of estrogens which primarily influence liver biology through the activation of Estrogen Receptor alpha (ERα). However, how ERα signaling in hepatocyte modulates liver functions according to the nutritional status, a critical determinant of hepatic metabolism, remains to be characterized.

The present study first reveals the nycthemeral expression profile of liver ERα, which peaks during the feeding period (ZT16) in both male and female mice. ERα expression is altered by reprogramming the hepatic circadian clock in response to inverted food intake, highlighting the influence of feeding on liver ERα expression patterns. As ERα is mainly expressed in hepatocytes both in human and mouse livers, the functional role of hepatocyte ERα during fasting or feeding state was then delineated, using mice harboring ERα hepatocyte-specific deletion (ERα^hep−/−^) compared with their wild-type littermates (ERα^hep+/+^). Transcriptomic analyses reveal significant sex differences in the adaptation of liver functions to this contrasted nutritional status, as well as the sex-specific regulatory effects of hepatocyte ERα. In such physiological settings, hepatocyte ERα deletion does not influence liver transcriptomic profiles in males but significantly alters gene expression in female livers, both fed and fasted conditions. Hepatocyte ERα more specifically controls genes associated with inflammatory responses and fatty acid metabolic pathways in females, modulating the ChREBP and PPARα pathways in feeding or fasting states, respectively.

Thus, hepatocyte ERα is integral to liver metabolic adaptations to fasting and feeding in females but not in males, thus exhibiting both sex-specific and diet-dependent actions. Further characterizing sex differences in liver metabolic flexibility, these results provide new information to develop sex-based strategies for the prevention and management of MASLD.

## INTRODUCTION

The liver has been recognized as one of the most sexually dimorphic organs (1–3). Supporting this assertion, extensive transcriptomic analyses conducted in both preclinical models and humans demonstrated that male and female livers exhibit distinct regulation of genes involved in xenobiotic metabolism, metabolic processes, and inflammatory responses (4–6). Notably, most studies have been conducted in pathophysiological settings, such as diet-induced obesity in animal models or human liver biopsies obtained from individuals with severe obesity and/or liver disease, thereby complicating the elucidation of authentic sex-specific hepatic functions in physiological contexts (7).

Sex chromosomes and endogenous sex steroid hormones, primarily androgens and estrogens, are the main determinants of these sex differences (8,9). Recent observations from mouse models that differentiate the effects of various sex-biasing factors on transcriptomic regulation in metabolic tissues have revealed that gonadal hormones exert a more pronounced influence than sex chromosomes on gene expression in the liver and adipose tissues (10). More specifically, estrogens play a significant role in whole-body metabolic homeostasis and are associated with metabolic advantages in humans, contributing to the lower susceptibility of premenopausal women to metabolic disorders such as type 2 diabetes and metabolic-associated steatotic liver disease (MASLD) compared to age-matched men (11,12). Decline in endogenous estrogen levels induced by spontaneous or surgical menopause (e.g., bilateral oophorectomy) promotes fat redistribution towards visceral depots and ectopic sites, including the liver, leading to metabolic dysfunctions that can be mitigated by estrogen-based hormonal therapies (13,14).

In murine models, estrogen deficiency, either due to genetic abrogation of E2 synthesis via inactivation of the Aromatase gene (Cyp19A1) or to bilateral oophorectomy, also results in adiposity, insulin resistance, and liver steatosis, conditions that can be prevented or reversed by 17β-estradiol (E2) administration (15,16). It is now well established that activation of the estrogen receptor alpha (ERα) mediates the metabolic actions of estrogens, including the prevention of liver steatosis (9,12,17,18). Moreover, recent reports provided evidence that ERα signaling in hepatocytes contributes to E2-mediated liver protection. Indeed, mice harboring hepatocyte-specific ERα deletion (known as LERKO or ERα^hep−/−^ mice) are more prone to high-fat diet-induced steatosis than their wild-type counterparts and no longer protected from liver lipid accumulation by E2 administration (15,19).

A comprehensive understanding of the physiological role of hepatocyte ERα in sex-specific liver metabolic regulation is essential to elucidate the mechanisms by which estrogens exert their protective action against MASLD pathogenesis. Previous studies demonstrated that ERα is more highly expressed in the liver of female mice as compared to males, and has a direct effect on the regulation of the hepatic genes relevant for energy metabolism. In females, the transcriptional activity of the receptor fluctuates with the phases of the estrous cycle, suggesting that ERα signaling is finely tuned to align liver functions with the energetic demands of each reproductive stage (20). Indeed, in response to nutritional intakes, and more specifically dietary amino acids, activation of hepatocyte ERα contributes to cholesterol and lipoprotein metabolism, but also regulates IGF1 production which is critical for uterine endothelium proliferation and progression of the estrous cycle (21,22). However, the specific role of hepatocyte ERα in the physiological adaptation of liver metabolism to varying nutritional states, namely fasting (a catabolic phase) and feeding (an anabolic phase), remains poorly understood.

Nutritional status significantly influences fundamental liver functions by engaging specific biochemical pathways that adapt whole-body energy homeostasis to available fuel sources. Therefore, the liver exhibits remarkable metabolic flexibility, a characteristic that may be subjected to sex differences and thus contribute to the lower susceptibility of females to MASLD. The objective of the present study was to elucidate the sex-specific influence of fasting and feeding phases on the expression level of hepatocyte ERα and its ability to regulate liver metabolic functions.

## RESULTS

### 1. Liver ERα gene expression is circadian

Circadian rhythms orchestrate crucial functions in the liver notably at the transcriptional level (23–26). Before deciphering the sex-dependent role of ERα in hepatic functions, we first examined the circadian expression profile of the receptor in the liver of male and female wild-type mice. Mice were fed *ad libitum* with a standard chow diet and harvested every 4 hours over a complete day and night cycle. To ensure controlled exposure to endogenous estrogens, females were synchronized in proestrus/estrus phase. Higher body weight and perigonadal white adipose tissue depots, as well as increased plasma levels of total and HDL-cholesterol, were observed in male compared to female mice while no sex differences were observed in other phenotypic characteristics including relative liver weight (**Fig. S1 A-H, left panels**). In the liver, a circadian variation was found in *Esr1* gene (encoding ERα) expression level in both sexes, rising at zeitgeber time (ZT) 16 during the dark activity/feeding period (ZT12-ZT24) while the nadir was detected during the light/resting period (ZT0-ZT12) (**Fig. 1A, left panel**). Of note, females exhibit higher receptor expression levels that males at every time point assessed. Other known estrogens receptors, i.e. ERβ (*Esr2*) and *Gpr30*, were detected at very low levels whatever the ZT and did not display such a nychthemeral expression profile in the liver (**Fig. 1B, left panels**). Androgen (*Ar)* and progesterone (*Pr)* receptor genes were weakly expressed as compared to *Esr1*, in contrast to glucocorticoid receptor (*Gr*) (**Fig. 1C, left panels**). However, none of them showed circadian expression. These results thus indicate that amongst the main steroid receptors, ERα is the only one that displays a rhythmic hepatic gene expression in both sexes.

**Figure 1.**
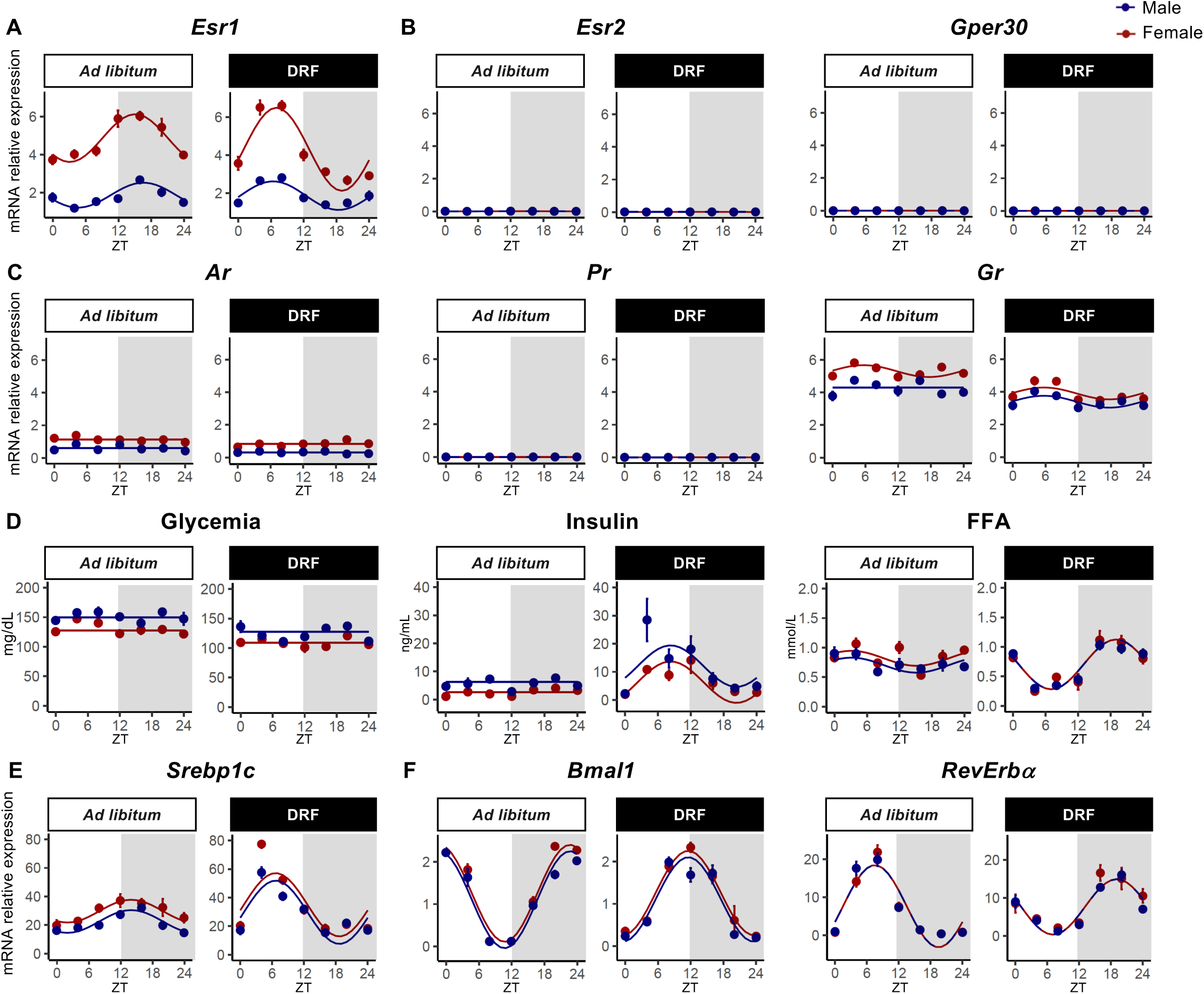
Liver ERα gene expression is circadian and controlled by food intake in male and female mice. Eleven-week-old male and synchronized female C57BL/6J mice were fed *ad libitum* or subjected to a day-restricted feeding (DRF, ZT0-ZT12) during 2 weeks (n=5 mice/sex/genotype/ZT). **(A-C)** Relative liver gene expression of ERα (*Esr1*) (A), ERβ (*Esr2*) and *Gper30* (B), *A*r, *Pr* and *Gr* (C) determined by qRT-PCR. **(D)** Quantification of circulating glucose, insulin and FFA levels in mice at the end of the experiment. **(E-F)** Relative liver gene expression of *Srebp1c* (E), *Bmal1* and *RevErbα* (F) determined by qRT-PCR. TBP was used as housekeeping gene. Data were analyzed with drylm function from the dryR package. Groups sharing the same fitting line shape have the same rhythmic parameters. A flat line signifies no rhythm detected.

### 2. Hepatic ERα gene rhythmicity is driven by food intake

Food intake is a powerful driver of the circadian clock expression and activity in the liver (27,28). To decipher whether changes in hepatic ERα expression are also dependent from feeding, we reprogrammed the liver clock by a daytime inverted restricted feeding (DRF) that uncouples the liver from the central clock (27). Male and synchronized female wild-type mice (*ERα^hep+/+^*) were adapted to eat only during the light phase (ZT0-ZT12) for 15 days. As compared to mice fed *ad libitum*, glucose, insulin and free fatty acids (FFA) plasma concentrations were consistent with the pattern of feeding behavior implemented in DRF experiments (**Fig. 1D**). Accordingly, mRNA expression profile of *Srebp1c*, known to be induced by food intake, as well as *Bmal1* and *RevErbα*, two key molecular actors of the circadian core clock, were reprogrammed (**Fig. 1E and F**). In contrast to other steroid hormone receptors, ERα gene expression profile in the liver was reprogrammed by DRF (**Fig. 1A-C, right panels**), demonstrating that food intake is a crucial driver of hepatic ERα gene expression.

### 3. Hepatic ERα is not required for the maintenance of the core clock gene expression

It has been suggested that ERα could regulate the circadian clock by itself (29,30). Thus, we then investigated whether ERα signaling in hepatocytes controls the expression of hepatic circadian core-clock and the clock-controlled genes. Mice with hepatocyte-specific ERα deletion (*ERα^hep−/−^*) and their control littermates (*ERα^hep+/+^*) were fed a standard chow diet either *ad libitum* (AL) or subjected to DRF. Regardless of sex, ERα deletion did not alter body weight, liver weight, perigonadal fat mass nor triglycerides and cholesterol circulating profiles (**Fig. S2)**. As expected, DRF modified plasma levels of glucose, insulin and FFA as compared to *ad libitum* feeding, with insulin concentrations being slightly lower in both male and female *ERα^hep−/−^* mice (**Fig. 2 A-C**). Hepatic expression of *Srebp1c* and *G6pc*, reflecting neoglucogenesis and *de novo* lipogenesis activity, respectively, was also effected by DRF (**Fig. 2D and E**). In both males and females, hepatocyte-specific *ERα* deletion did not alter the circadian gene expression profile of the core loop (Clock, Bmal1, Cry 1/2 and Per 1/2) (**Fig. 2F**), the interlocking loop (Rev-Erbα) (**Fig. 2G**), nor the expression of output genes (Nfil3, Tef and Hlf) (**Fig. 2H**).

**Figure 2.**
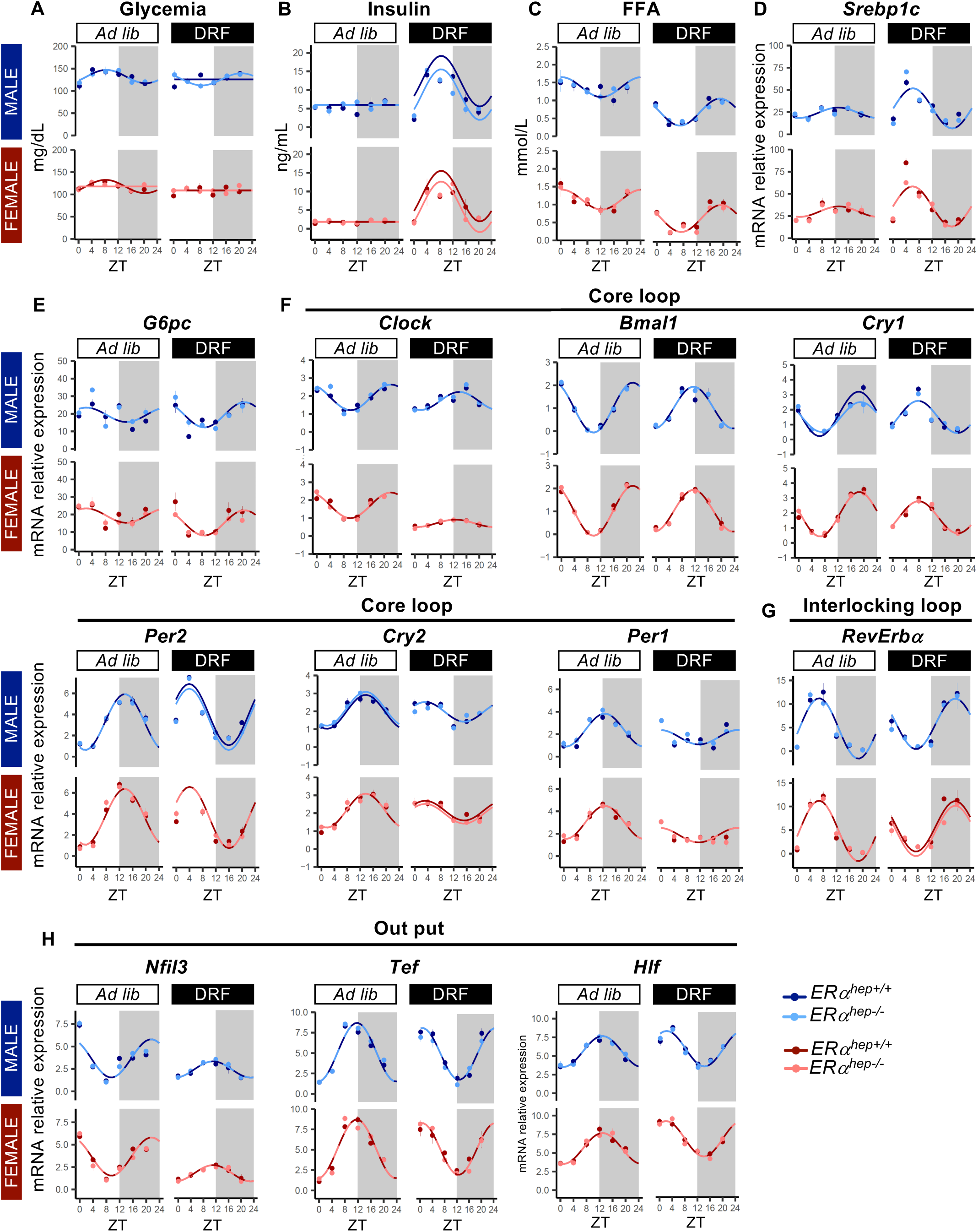
Liver circadian clock is not influenced by hepatocyte ERα deletion in both sexes. Eleven-week-old male and synchronized female hepatocyte-specific knockout mice (*ERα^hep−/−^*) and their control littermates (*ERα^hep+/+^*) were fed *ad libitum* or subjected to a day-restricted feeding (DRF, ZT0-ZT12) during 2 weeks (n=5 mice/sex/genotype/ZT). **(A-C)** Quantification of circulating glucose (A), insulin (B) and FFA (C) levels in mice at the end of the experiment. **(D-E)** Relative liver gene expression of *Srebp1c* (D) and *G6pc* (E) determined by qRT-PCR. **(F-H)** Relative liver gene expression of liver clock genes *Clock*, *Bmal1*, *Cry1*, *Per2*, *Cry2* and *Per1* (F), *RevErbα* (G), *Nfil3*, *Tef* and *Hlf* (H) determined by qRT-PCR. TBP was used as housekeeping gene. Data were analyzed with drylm function from the dryR package. Groups sharing the same fitting line shape have the same rhythmic parameters. A flat line signifies no rhythm detected.

Altogether, our data indicates that, in both sexes, the circadian liver expression of *ERα* is regulated by food intake whilst ERα signaling in hepatocyte is not involved in the regulation of the liver core-clock gene expression.

### 4. Hepatocytes mediate feeding-dependent regulation of ERα expression

We then sought to determine which cell types express ERα within the liver and, among them, which ones present a rhythmic expression of the receptor. Using available transcriptomic databases obtained from Human (31) or mouse (Tabula Muris, https://tabula-muris.ds.czbiohub.org/) liver samples, we first determined that, in both species and both sexes, *Esr1* expression is mainly confined to hepatocytes and, to a lesser extent, in sinusoidal endothelial cells (**Fig. 3A and B**). Results obtained from male and synchronized female *ERα^hep+/+^* and *ERα^hep−/−^* mice, either fed *ad libitum* or subjected to a DRF, confirmed that *Esr1* expression in the liver primarily occurs in hepatocytes and suggested that the rhythmic profile observed in *ERα^hep+/+^* control mice mainly reflects hepatocyte expression (**Fig. 3C**). To further investigate the influence of the nutritional status on hepatocyte ERα expression, male and synchronized female *ERα^hep+/+^* and *ERα^hep−/−^* mice were harvested at ZT16 (time characterized by the higher level of *ERα* expression in mice fed *ad libitum*) either fed *ad libitum* or fasted for 16 hours (from ZT0 to ZT16). The experimental design, aiming to compare feeding to fasting status, was validated by the patterns of plasma glucose, insulin and FFA, as well as by changes in gene expression of G6pc and Srebp1c (**Fig. S3 A-E**). Consistent with the circadian profiles in **Fig. 1A and Fig. 3C**, hepatic ERα mRNA and protein levels were enhanced by feeding in both sexes, while expression remained higher in females (**Fig. 3D**). Of note, bilateral oophorectomy did not affect this feeding-related regulation in female mice (**Fig. 3E**).

**Figure 3.**
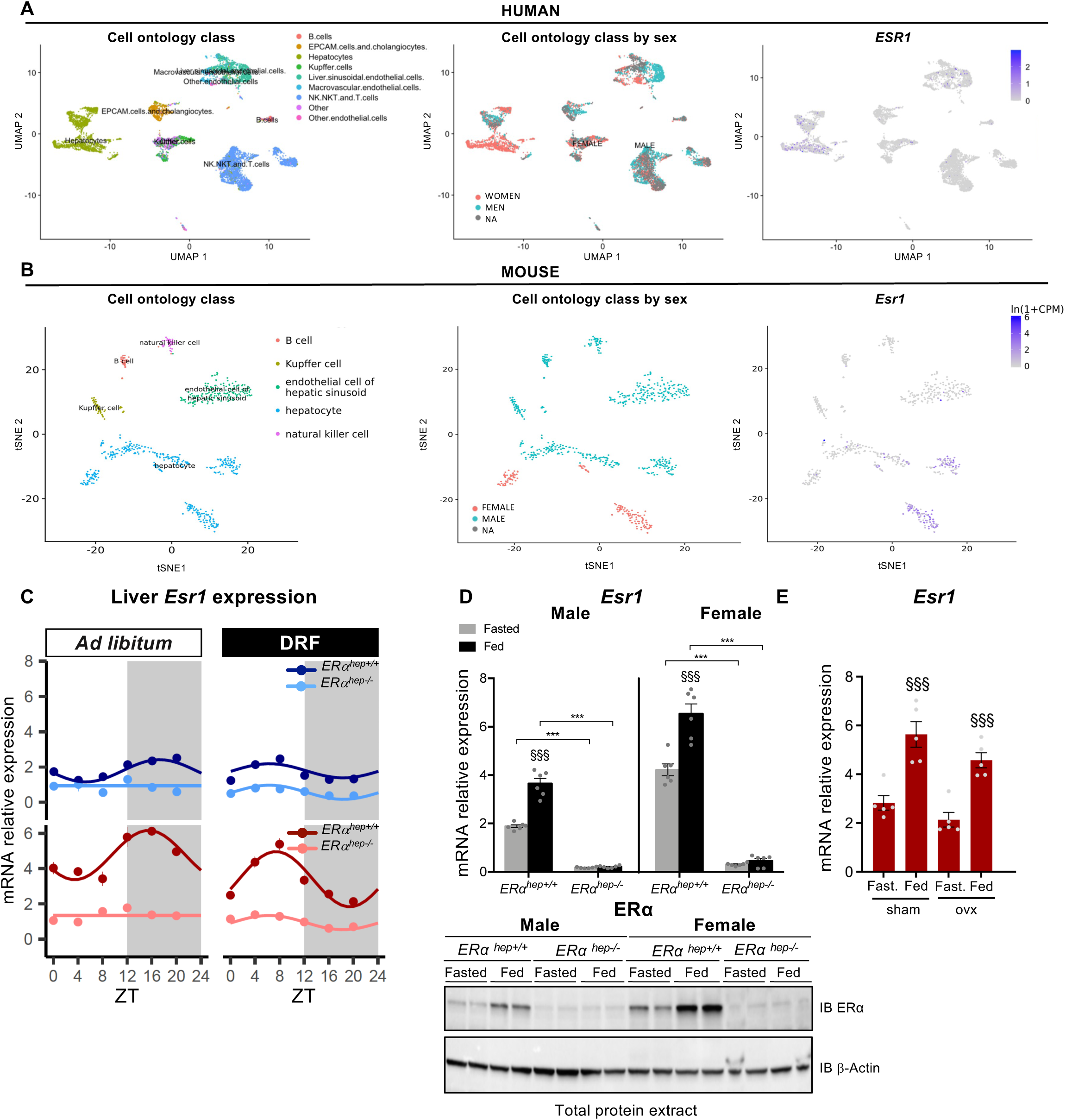
Liver ERα is mainly expressed in hepatocyte in Human and mice. **(A-B)** Cell ontology class by liver cell types, sex and expression of ERα in Human (ESR1, A) and mice (Esr1, B) from ScRNA-Seq human dataset (31) and murine database (Tabula Muris) analysis. **(C)** Eleven-week-old male and synchronized female hepatocyte-specific knockout mice for ERα (*ERα^hep−/−^*) and their littermate controls (*ERα^hep+/+^*) were fed *ad libitum* or subjected to a day-restricted feeding (DRF, ZT0-ZT12) during 2 weeks (n=5 mice/sex/genotype/ZT). Relative liver mRNA expression of *Esr1* (ERα) determined by qRT-PCR. TBP was used as housekeeping gene. Data were analyzed with drylm function from the dryR package. Groups sharing the same fitting line shape have the same rhythmic parameters. A flat line signifies no rhythm detected. **(D)** Eleven-week-old male and female *ERα^hep−/−^* and *ERα^hep+/+^* mice were fed *ad libitum* or fasted from ZT0, then euthanized at ZT16 (n= 6 animals/group). (D) Relative liver mRNA expression of *Esr1* (ERα) determined by qRT-PCR. TBP was used as housekeeping gene. Liver *ERα* protein expression was determined by Western Blot (n=2 animals/experimental condition). **(E)** *ERα^hep+/+^* mice were ovariectomized at the age of 8 weeks and then euthanized at ZT16 fed ad libitum or fasted for 16 hours at 11-week-old. Relative liver mRNA expression of *Esr1* (ERα) determined by qRT-PCR. TBP was used as housekeeping gene. Data represent mean ± SEM; statistical significance was calculated using two-way ANOVA with Sidak’s multiple comparisons. §, nutritional effect; *, genotype effect. §§§ or ***, p-value < 0.001.

Altogether, our findings demonstrate that ERα expression in the liver follows a rhythmic expression pattern and is mainly expressed in hepatocytes in both males and females. Moreover, our results suggest that, rather than estrogen availability, nutritional status serves as the primary regulator of hepatic ERα expression.

### 5. Hepatic transcriptomic adaptations to fasting and feeding are sex-specific

We then investigated sex-specific hepatic adaptations to feeding and fasting periods, a physiological aspect that remains poorly explored so far. Liver gene expression profiles were analyzed by microarrays in male and synchronized female *ERα^hep+/+^* mice submitted to contrasted physiological nutritional conditions i.e. fed *ad libitum* or fasted, as previously described (**Fig. 3D)**. Large-scale transcriptomic profiles first revealed a clear influence of sex on liver adaptation, as illustrated on principal component analysis (PCA) which discriminated both sex and nutritional status (**Fig. 4A**). Highlighting the genes driving the separation of the four groups in the PCA, the upset plot (**Fig. 4B**) identified 1,083 genes associated with Dim2 and 113 genes with Dim1, primarily reflecting sex- and nutrition-related differences, respectively. Additionally, 70 genes were influenced by both sex and nutritional status. Gene Set Enrichment Analysis (GSEA) from the upset plot revealed that immune functions discriminated males from females (**Table S1**) and translation process, fatty acid and carbohydrate metabolic processes discriminated the nutritional effect (**Table S2**). We then performed a hierarchical clustering of the top 5,700 gene probes (4,741 differentially expressed genes, DEG), selected with an adj. p value of 0.05 and a fold change threshold of 1.5, that differ between livers of fasted and fed wild-type *ERα^hep+/+^* males and females. Eight broad gene clusters and their mean z-score were identified (**Fig. 4C and D**). Clusters 1 and 7 consisted of genes whose expression was unaffected by nutritional status but exhibited marked sex-dependent differences, with higher expression in males (cluster 1) or females (cluster 7), respectively. Cluster 1 included genes involved in xenobiotic metabolism as well as arachidonic and linoleic acid metabolic pathways, predominantly regulated by the transcription factors SF1 (Nr5a1) and Hnf4α (**Fig. 4E and F**). In contrast, genes in cluster 7 were associated with sulfation processes and immune-related functions under the control of Spi1, Irf8, AhR and of the nuclear receptors PXR (Nr1i2) and SF1 (Nr5a1). Cluster 6 was composed of genes significantly upregulated during fasting in both sexes, enriched for pathways related to fatty acid oxidation. These genes are submitted to the transcriptional regulation of PPARα, Creb1 and Foxo1, three well-established drivers of the fasting response (32,33). In contrast, clusters 3 and 4 included genes upregulated in the fed state across sexes, but that exhibited sex-specific repression during fasting with downregulation more pronounced in females (cluster 3) or in males (cluster 4). Notably, despite fasting-induced repression, genes in cluster 4 remained more highly expressed in females than in males under fasting, whereas cluster 3 shown an opposite trend. Functional enrichment analysis indicated that cluster 3 was associated with glucose-responsive pathways, including those regulated by ChREBP, while cluster 4 was enriched for genes involved in sterol, cholesterol, and lipid biosynthesis, as well as steroid metabolism, governed by SREBP1c, SREBP2, and the nuclear receptors COUP-TFII (Nr2f2) and ERα (Esr1). Finally, clusters 2, 5, and 8 -linked to vasculogenesis and angiogenesis (cluster 2), DNA replication (cluster 5), and immune responses (cluster 8) -were hugely regulated by nutritional status in females but only modestly in males. These responses involve key transcriptional regulators such as NF-κb, Spi1, or Ets1. Interestingly, the hepatic associated gene correlation network analyses performed for PPARα (cluster 6), ChREBP (cluster 3) and ERα (cluster 4), that came out to be sex-specific or not, revealed a different pattern of co-correlated genes in females and males, suggesting that their respective activities were sexually dimorphic (**Fig. 4G**).

**Figure 4.**
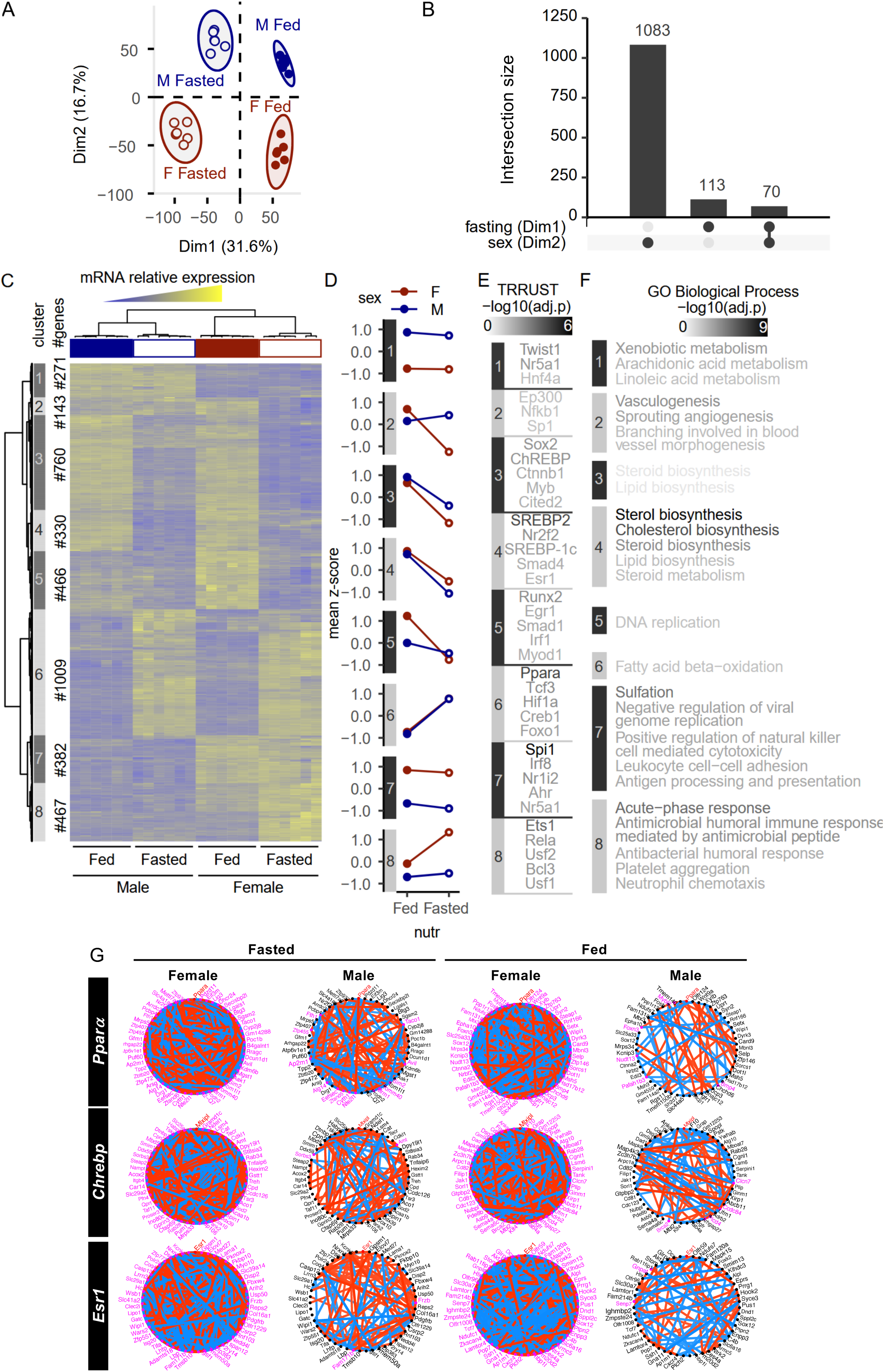
Sex-specific influence of hepatocyte ERα on liver transcriptome in fasting and feeding conditions. Twelve-week-old male and synchronized female hepatocyte-specific knockout mice for ERα (*ERα^hep−/−^*) and their littermate controls (*ERα^hep+/+^*) were fed *ad libitum* or fasted for 16h and then euthanized at ZT16 (n=6/group). Liver transcriptome was determined by microarrays. **(A)** Principal component analysis of liver transcriptome of fed and fasted male and female *ERα^hep+/+^* mice. **(B)** Upset plot established from PCA. **(C-D)** Heatmap (C) representing DEG (p-value<0.05, |FC|>1.5) from male and female *ERα^hep+/+^* liver transcriptome and cluster profile (D). **(E-F)** Transcriptional factors involved in these pathways were listed using TTRUST database (E) and Gene Ontology analysis performed on each cluster (F). **(G)** Networks of the 50 first genes showing the highest absolute correlation with *Pparα*, *Chrebp* and ERα (*Esr1*) in fasting and feeding. The edges corresponding to significant correlations are presented (Bonferroni-adjusted p<0.05). For each nutritional status, the network circle plot based on the top 50 genes selected in female and the corresponding network in males are presented on the left and on the right, respectively. Magenta nodes correspond to genes that significantly correlate with the gene of interest. Red and blue were used for positive and negative correlations, respectively.

Altogether, our data show that livers of male and female mice share fundamental biological functions, allowing them to adapt to the appropriate nutritional status. They also displayed sex-specific gene expression patterns and gene correlation networks according to fasting or feeding condition.

### 6. Sex-specific role of hepatocyte ERα in liver adaptation to nutritional status

Analysis of our own transcriptomic database from human and mouse liver samples (GSE159090, GSE159088, respectively) (34) shown that ERα co-regulated gene networks were significantly different in males and females, suggesting sex-specific functions for liver ERα (**Fig. S4**). While hepatocyte-specific ERα deletion did not impact male liver transcriptome whatever the nutritional setting, it strongly altered female liver transcriptomic signature in both feeding and fasting conditions (**Fig. 5A and Fig. S5**). To identify the genes driving the PCA-based segregation, the upset plot (**Fig. 5B**) revealed that 1,415 and 932 genes contribute to the sex effect (Dim2) and the nutritional effect (Dim 1), respectively. Notably, 1,375 genes (Dim 3) accounted for the genotype effect specifically in females and 22 genes were jointly influenced by sex and genotype. These findings highlighted the sex-specific role of hepatocyte ERα in shaping the female liver transcriptome, an observation further supported by the hierarchical clustering analysis (**Fig. 5C**). Performed on the 1363 dysregulated genes in females *ERα^hep−/−^* vs *ERα^hep+/+^* in both nutritional status, the analysis identified four distinct clusters. Clusters 1 and 4 contained genes whose expression was not affected in male whatever the nutritional condition while they were modulated in control *ERα^hep+/+^* females and affected by ERα deletion in hepatocyte (**Fig. 5D**). Cluster 1 genes were increased by fasting in female compared to male *ERα^hep+/+^* mice and had an inverted expression pattern in hepatocyte-ERα deleted mice. They are related to extracellular matrix (fibrinolysis, cell-cell adhesion or extracellular matrix organization), acute-phase and adaptative immune responses, controlled notably by Stat1, HNF4α, ERα (Esr1), Snail and SHP (Nr0b2) (**Fig. 5E and F**). Cluster 4 gene expression was higher in females from both genotypes compared to males and hepatocyte ERα deletion abolished their fasting-induced downregulation. These genes are associated with GO functions involved in morphogenesis, and vasculogenesis, and their regulation involves the transcription factors Runx2, Twist1 and Ctnnb1. Clusters 2 and 3 contained genes whose expression pattern is inversely regulated by the nutritional status in male and female *ERα^hep+/+^* mice, i.e. up or down-regulated in fasted *ERα^hep+/+^* females while down- or up-regulated in *ERα^hep+/+^* males, respectively. Hepatocyte-specific ERα deletion led to a masculinization of their expression pattern, i.e. down and up-regulated in fasting compared to feeding for clusters 2 and 3, respectively. Cluster 2 encompassed genes related to humoral immune response and cluster 3 genes were associated with GO functions involved in retinol metabolism, fatty acid and acyl-coA metabolism, tricarboxylic acid cycle (TCA), vasculogenesis and canonical Wnt and SMAD signaling pathways involving the transcription factors Myc, Sp1 and Trp53.

**Figure 5.**
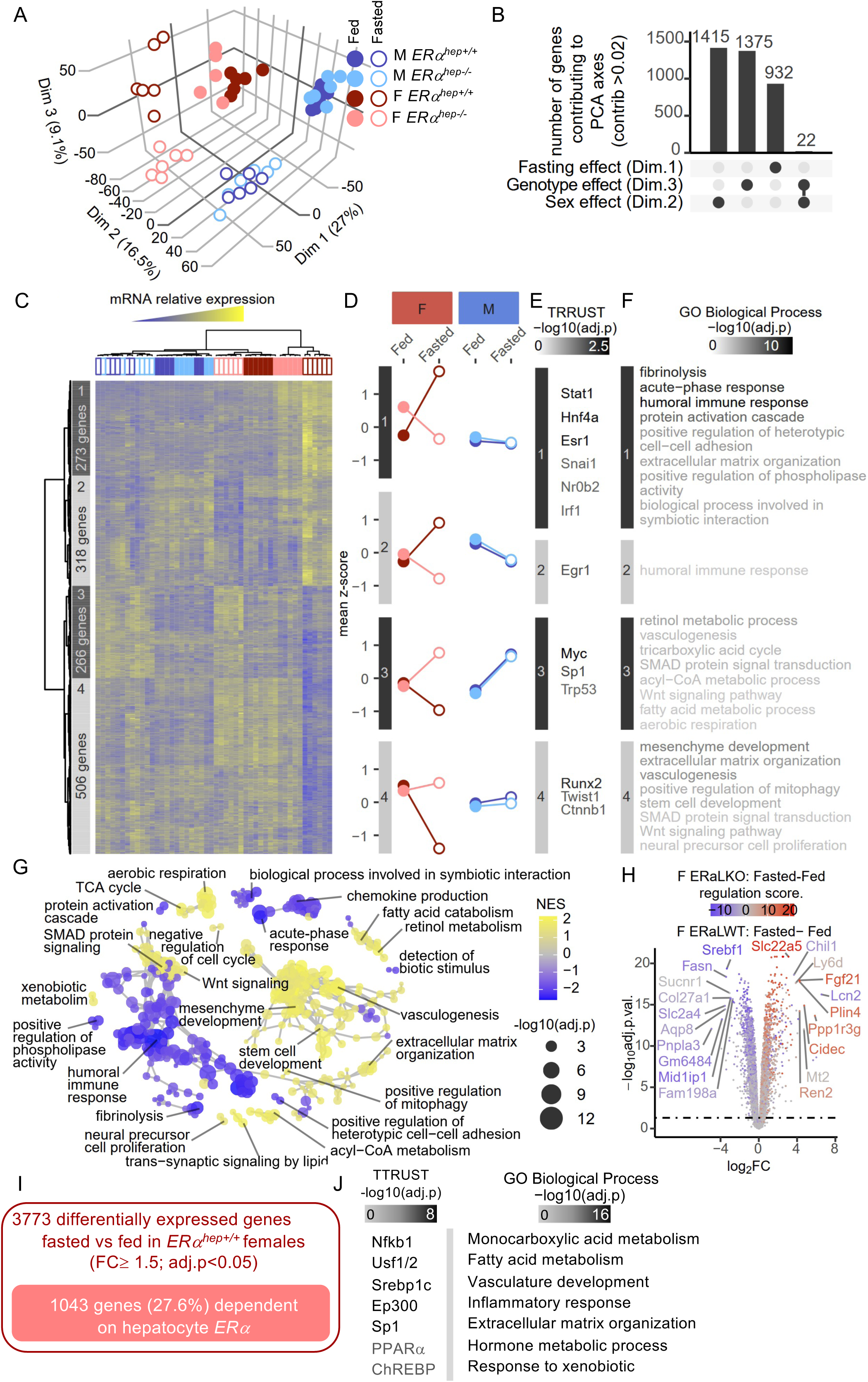
Sex differences in hepatocyte ERα functions in male and female mice. Twelve-week-old male and synchronized female hepatocyte-specific knockout mice for ERα (*ERα^hep−/−^*) and their control littermates (*ERα^hep+/+^*) were fed *ad libitum* or fasted for 16h and then euthanized at ZT16 (n=6/group). Liver transcriptome was analyzed by microarrays. **(A)** Principal component analysis of liver transcriptome in male and female *ERα^hep+/+^* and *ERα^hep−/−^* mice. Each dot represents an observation (animal), projected onto three dimensions: first dimension, nutritional effect; second dimension, sex effect; third dimension, genotype effect. **(B)** Upset plot illustrating the number of genes contributing to PCA axis. **(C-F)** Hierarchical clustering (C) representing DEG (p-value<0.05, |FC|>1.5) in male and female *ERα^hep+/+^* and *ERα^hep−/−^* liver transcriptome and cluster profile (D). Transcription factors involved in each cluster were defined using TTRUST database (E) and Gene Ontology analysis was performed (F). **(G)** Gene Set Enrichment Analysis performed on differentially expressed genes (p-value<0.05, |FC|>1.5) between fed and fasted *ERα^hep+/+^* and *ERα^hep−/−^* female mice. **(H)** Volcano plot illustrating *ERα^hep−/−^* effect on gene expressed in fasted and fed *ERα^hep+/+^*. **(I-J)** Hepatocyte ERα-dependent impact on liver expressed genes in fasted and fed *ERα^hep+/+^* female (p-value<0.05, |FC|>1,5) in females and transcription factors involved defined using TTRUST database and Gene Ontology analysis (J).

Focusing on the effect of hepatocyte ERα deletion in females, Gene Set Enrichment Analysis (GSEA) from the genotype effect (Dim 3) of the upset plot (**Fig. 5B**) further highlights the results of the hierarchical clustering. Interestingly, amongst the more regulated hepatic genes in fasted and feeding *ERα^hep+/+^* females, hepatocyte ERa deletion impacted the expression of Srebf1 (Srebp1c), Fasn, Pnpla3, Plin4, Cidec that are all involved in intracellular FA homeostasis and on the expression of the hepatokine Fgf21 (**Fig. 5H**). We then sought to question the influence of hepatocyte ERα on the female-specific liver trancriptome and the regulation of sex-biased genes (SBG). Data showed that 27.6% of the DEG in fasted *vs* fed condition in *ERα^hep+/+^* females were dependent of ERα signaling in hepatocyte (**Fig. 5I and Tables S3 and S4**), representing 19.7% of the SBG (**Fig. S7 and Tables S5 and S6**). GO analysis indicated that *ERα* hepatocyte-specific deletion alters biological functions related to fatty acid metabolism, vasculature development and inflammatory responses involving, notably, the transcription factors Nf-kb1, Usf1/2, Ep300, Sp1, Srebp1c, ChREBP, and PPARα (**Fig. 5I and Table S4**). Interestingly, hepatocyte ERα-dependent transcriptomic signatures revealed overlaps with sex-specific biological processes and transcription factors previously identified as different between males and females but also common to both sexes (**Fig. 4D-F**). Indeed, hepatocyte ERα deletion affected not only biological processes specific to female mice (namely inflammatory and acute-phase responses or vasculature development), but also processes shared by both sexes, related to fatty acid metabolism involving the transcription factors PPARα, Srebp1c and ChREBP.

Altogether, the results highlight the role of ERα in female liver physiology by modulating sex-specific and common biological functions, especially fatty acid metabolism.

### 7. Hepatocyte ERα regulates liver fatty acid metabolism in a sex-specific manner

The transcriptomic data emphasized a role for hepatocyte ERα in lipid metabolism involving the transcription factors Srebp1c, ChREBP and PPARα, which are activated by feeding (involving insulin and glucose signaling pathways) and fasting, respectively (**Fig. 5J**). To further investigate this fundamental liver function, we first used a consensus genome-scale metabolic model called iHepatocytes (35) to predict metabolites that could be affected by ERα hepatocyte deletion. The resulting metabolic network highlighted the metabolites that were predicted to be significantly different in the absence of hepatocyte ERα (**Table S4**). In agreement with this analysis, ChREBP gene expression and activity, which was reflected by *Lpk*, *Elovl6*, were specifically downregulated in the liver of fed *ERα^hep−/−^* female mice compared to their control littermates (**Fig. 6A-B**). No effect was observed on *Srebp1c* expression (data not shown). However, *Fasn*, whose gene expression is induced by both ChREBP and Srebp1c, was decreased in liver of *ERα^hep−/−^* females (**Fig. 6C**). In fasting condition, PPARα expression and activity, reflected by its target genes *Cyp4a14* and *Vnn1*, were specifically upregulated in *ERα^hep−/−^* female mice (**Fig. 6D-E**). Accordingly, fasting-induced increase in β-hydroxybutyrate plasma levels was more pronounced in *ERα^hep−/−^* female mice than in their control littermates, while fasted glycemia, respiratory exchange rate (RER), energy expenditure, activity or food intake were not affected (**Fig. 6F and Fig. S8**).

**Figure 6.**
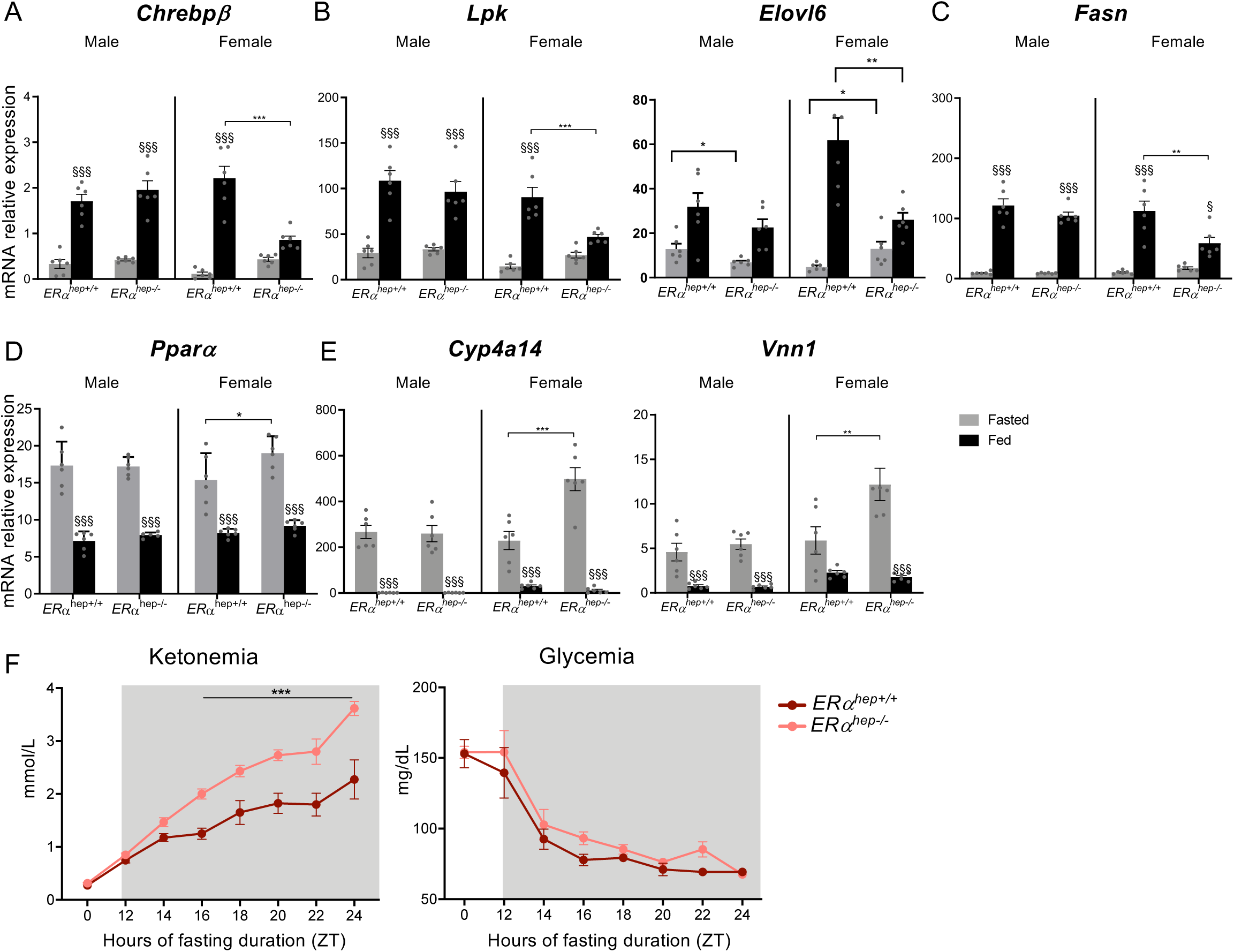
Hepatocyte ERα controls fatty acid lipid metabolism in female mice. **(A-E)** Twelve-week-old male and synchronized female hepatocyte-specific knockout mice for ERα (*ERα^hep−/−^*) and their control littermates (*ERα^hep+/+^*) were fed *ad libitum* or fasted for 16h and then euthanized at ZT16 (n=6/group). Relative liver mRNA expression of genes involved in *de novo* lipogenesis (B-E) and in fatty acid oxidation (fasting) (F-H) determined by qRT-PCR. TBP was used as housekeeping gene. Data represent mean ± SEM; statistical significance was calculated using two-way ANOVA with Sidak’s multiple comparisons. §, nutritional effect; *, genotype effect. § or *, p-value < 0.05; §§ or **, p-value < 0.01; §§§ or ***, p-value < 0.001. **(F)** Twelve weeks-old synchronized female ERα hepatocyte-specific deleted mice (*ERα^hep−/−^*) and their control littermates (*ERα^hep+/+^*) were fasted for 24 hours (from ZT0 to ZT24, n=5/group). Glycemia and ketonemia were measured at ZT0 and then every 2 hours from ZT12 to ZT24. Repeated measures ANOVA were used to compare changes over time between the genotypes. *, genotype effect, *** p-value < 0.001.

Demonstrating the modulation of master regulators of liver metabolism in feeding and fasting states, these data highlight the crucial contribution of hepatocyte ERα to lipid metabolism homeostasis in liver of female mice. They also emphasize a nutritional-driven function for hepatocyte ERα and the need to systematically consider the nutritional status to explore the precise influence of the receptor on liver functions.

## DISCUSSION

The present study provides further evidence that the liver is one of the most sexually dimorphic organ, as previously demonstrated by the differential expression between males and females of hundreds of genes that govern specific biological functions (1,2,6). However, so far, no study deeply investigated sex specificities in the adaptation of liver functions to fasting and feeding periods under physiological conditions. Our experimental approaches, applied to mice fed a standard chow diet and synchronized in pro-estrus/estrus for female mice, have made it possible to characterize sex-specific transcriptomic signatures associated with controlled hormonal status and contrasted nutritional status, then to determine the selective contribution of hepatocyte ERα to the modulation of gene expression in female liver.

Sex differences in liver functions primarily result from the actions of sex steroid hormones, over the influence of sex chromosomes (10). More specifically, estrogens are now recognized to confer liver protection against nutritional challenges promoting the development of MASLD, through the activation of ERα (11,12). Our results confirm that, in contrast to other estrogen receptors, ERα is highly expressed in the liver of mice of both sexes but with significantly higher mRNA and protein levels detected in female than in male mice (**Fig. 1 and Fig. 3**). Moreover, among receptors for steroid hormones, ERα is the only one to display a rhythmic expression over the clock with a nadir at ZT16, during the feeding/activity period (**Fig. 1**). Although the nychthemeral expression of ERα has already been reported in male mice (36), the present study is the first to demonstrate such a regulation in females which exhibit a greater expression amplitude than males.

As in the majority of peripheral metabolic organs, it is well accepted that the main time giver (or Zeitgeber) for the liver circadian clock is represented by feeding cycles (37,38), underpinning a tightly crosstalk between dietary intake and the circadian timing system (27). Of note, submitting mice to a DRF, that reprograms the liver clock, reverses ERα circadian expression profile in both sexes (**Fig. 1A**), demonstrating the crucial role of food intake and/or the liver clock in the regulation of ERα expression. Whilst a relationship has already been described between estrogen signaling and circadian regulation in female rodents, notably with the central circadian timing of the hypothalamic-pituitary-ovarian axis (39,40), no impact of hepatocyte ERα was found here on the expression of the core-clock genes in male and female mice fed a standard chow diet (**Fig. 2**). This contrast with previous *in vitro* studies conducted on cancer cell lines that suggested a role for ERα in *Per2* and *Clock* expression (41,42). However, we cannot exclude a function for estrogens and, potentially for hepatocyte ERα, in the modulation of the liver clock under obesogenic conditions. In line with this hypothesis, some studies suggested an influence of estrogens on the liver clock when female mice were challenged by a long-term exposure to a HFD. In response to such a nutritional stress, Per2:Luc rhythm was advanced 4h in the liver of oophorectomized versus intact female mice (43), and the phase of the liver circadian clock was restored to normal when oophorectomized females were treated with E2 (44). Thus, estrogens seem to markedly alter the liver circadian clock under metabolic stress but whether this engages hepatocyte ERα requires further investigations. To our best knowledge, the present study is the first to report *in vivo* evidence of a feeding-driven, hepatic circadian regulation of ERα expression in both female and male mice. Besides the proper effect of the estrous cycle and associated variations in sex steroid hormone concentrations, evidence of a nutritional daily regulation of ERα expression brings a novel comprehensive level in the biology of this nuclear receptor. Indeed, leading to the conclusion that hepatocyte ERα expression is modulated by food intake, our findings also underscore time-dependent and diet-driven functions of ERα in the liver.

Supporting this hypothesis, our liver transcriptomic data suggest a close relationship between food availability and sex-specific ERα hepatocyte functions. In mice maintained on a standard chow diet, we found that hepatocyte ERα contributes to the adaptation of gene expression to fasting and feeding in the liver of female mice. In this physiological context, hepatocyte ERα mainly regulates the expression of genes involved in lipid metabolism, especially fatty acid metabolism, and inflammation, notably of acute phase responses (**Fig. 4**). Importantly, the liver is also viewed as an immunological organ, being responsible for the production of acute phase response proteins, complement components, cytokines and chemokines, and contains diverse populations of resident immune cells (45). This physiological property can be explained by its direct exposition to dietary and gut-bacterial products with inflammatory potential that requires a tight regulation of the inflammatory response to maintain tissue and organ homeostasis and to redistribute the energy resources during the raising of an inflammatory response (46–48). Inflammatory-associated biological processes are particularly sex-dependent as genes involved in these biological functions were more expressed in the liver of female compared to male control mice (**Fig. 3**). Numerous studies have described the close interplay between ERα-mediated actions of estrogens and anti-inflammatory processes in diverse immune cell types and organs, and in distinct pathophysiological contexts. Our findings showed that most of these inflammatory pathways were dependent on the nutritional status. Moreover, hepatocyte-specific ERα deletion leads to a misadaptation in the expression of genes encoding inflammatory effectors to the proper nutritional signal (**Fig. 4**), emphasizing the role of hepatocyte ERα in the coordination between metabolic regulations and immune responses. This is particularly the case for acute phase response genes whose expression and regulation profiles were inverted in the liver of in *ERα^hep−/−^* compared to *ERα^hep+/+^* female mice (**Fig. 4**). Whether this ERα function is due to the modulation of Nfkb1 or Stat activity, as described elsewhere (49,50), remains to be determined, as well as the involvement of hepatocyte ERα in chronic inflammatory liver diseases associated with metabolic disorders such as MASLD.

Besides inflammation, hepatocyte ERα is demonstrated here to be involved in fatty acid homeostasis by regulating their synthesis via *de novo lipogenesis* (DNL) and their degradation through β-oxidation, two crucial liver biological processes controlled upstream by the nutritional status (51,52). Indeed, hepatocyte ERα plays a positive role in DNL during the feeding/dark period by up-regulating ChREBP expression and activity, which was reflected by *Lpk* expression, a specific ChREBP target gene, and the expression of *Fasn* and *Elovl6* that were decreased in *ERα^hep−/−^* fed female mice compared to their control littermates (**Fig. 6**). During the light/resting period, hepatocyte ERα negatively controls fatty acid β-oxidation by down-regulating PPARα gene expression and transcriptional activity, reflected by the expression profile of its well-known target genes *Cyp4a14* and *Vnn1* (32) (**Fig. 6**). Accordingly, while the neoglucogenesis pathway was no impacted, fasting-induced plasma ketone bodies level was more elevated in *ERα^hep−/−^* compared to *ERα^hep+/+^* female mice.

Previous reports concluded that hepatocyte ERα does not mediate PPARα expression or activity in physiological condition but rather regulates cholesterol and lipoprotein metabolism in cooperation with LXRα (22,53). This discrepancy with our findings could be due to differences in the fasting duration and ZT used between the experimental settings. Indeed, using the PPARα hepatocyte-specific knockout mouse model, we recently demonstrated that a 16-hours fasting (from ZT0 to ZT16), as performed in the present study, is the optimal condition to properly study PPARα function (32). Of note, sex differences in liver PPARα biology has already been reported (34,54) and ERα binding sites have been described in *Pparα* promoter in the cardiac and skeletal muscle (55), as well as in breast cancer cells (56). However, the molecular mechanism mediating ERα-related regulation of PPARα gene expression and activity in the liver requires further investigations.

Some studies also described an interplay between ERα and ChREBP signaling. Indeed, E2-activated ERα has been shown to suppress lipogenesis by inhibiting mRNA and protein expressions of ChREBP and Srebp1c in insulin-secreting INS-1 cells (57). At the post-translational level, membrane ERα signalling inhibits triglyceride synthesis through the suppression of ChREBPα nuclear translocation *in vivo* in white adipose tissue and *in vitro* in adipocytes cell lines (58). More recently, using a hepatoma cell line, an interaction between ERα and ChREBP has been reported in the nucleus, leading to a decrease in ChREBPα-induced ChREBPβ transcription and promoting ChREBPβ degradation (59). Nevertheless, our analysis of ChIP-Seq experiment using anti-ERα antibody on liver samples of oophorectomized female mice treated or not with E2 (60) allowed us to identify ERα binding sites in the active, H3K27ac marked, promoter region of *Mlxipl* but not *Lpk* (data not shown). We also identified ERα binding sites in the active promoter/enhancer of *Fasn* and *Elovl6*, whose expression involved ChREBP and other key lipogenic transcription factors such as Srebp1c and LXRα (data not shown). This suggests that those genes could also be direct target genes for ERα. Those studies, performed in other tissues or cell lines than healthy liver or primary hepatocyte cultures, contrast with our findings that demonstrate a positive role of feeding-induced transcription factors on hepatocyte ERα expression and activity. However, we could speculate that, during feeding, hepatocyte ERα transcriptionally activates ChREBPα gene expression, which, in turn and together with ERα signaling, induces ChREBPβ expression. Then, in cooperation with ChREBP, Srebp1c and LXRα, hepatocyte ERα could participate to the expression of genes of the DNL pathway such as Fasn and Elovl6. However, the interaction between hepatocyte ERα and ChREBP or LXRα remains to be further demonstrated in our experimental conditions.

Altogether, the present study demonstrates that hepatocyte ERα biology is controlled by food availability, which contributes to the hepatic circadian expression profile of the receptor and drives its modulatory function on liver transcriptome. By integrating multiple input signals that reflect the metabolic status of hepatocytes, ERα should be considered as a metabolic sensor able to optimize liver adaptation to fasting and feeding conditions in female mice.

## Acknowledgements

We thank all members of the CREFRE staff, the GeT-Trix Genotoul facility, and Anexplo facilities for their help. B.T. was supported by FRM (FDM201906008682), SFD-Lilly and ANR Hepatomorphic (ANR-20-CE14-0035). This work was funded by ANR Hepatomorphic (P.G., A.M., C.P., H.D., H.G), FRM Metabotox, grant number ENV202109013962 (to H.G. with A.M. and P.G.), SFN (A.M.), SFD (A.M.) and by Région Occitanie (to N.L. with H.G., P.G., A.M.).

## Materials and Methods

### 1. Animals

Mice were housed in groups of 5 or 6 in polycarbonate cages and kept in a specific pathogen-free and temperature-controlled facility (22°C ± 2°C) on a 12-h light/dark cycle (ZT0-ZT12/ZT12-ZT24). ZT stands for Zeitgeber time, knowing that ZT0 is defined as the time when the lights are turned on. Six-week-old C57BL/6J male and female mice were purchased from Janvier Labs (Janvier Labs, Le Genest-Saint-Isle, France). Male and female mice harboring a specific deletion of *ERα* in hepatocytes were generated in the laboratory on a C57BL/6J genetic background by crossing mice floxed on exon 2 of *ERα* (*ERα ^flox/flox^*) with mice expressing Cre Recombinase under the control of the Albumin promoter (*ERα ^hep−/−^*). Wild-type littermate *ERα*-floxed mice (*ERα^hep+/+^*) were used as controls. The genotype of the mice was determined by PCR. At the age of 8 weeks, female mice were anesthetized with an intraperitoneal injection of ketamine (10 mg/kg) and xylazine (1 mg/kg) for ovariectomy. For the other experiments, female mice were synchronized at the proestrus/estrus (physiological peak of plasma estrogen concentration) stage of their estrous cycle at time ZT12 of the day of the experiment. Briefly, litter from a wild C57BL/6J male, sexually experimented, was placed in cages of female mice 62 to 64 hours before each experiment (protocol provided by Dr. Vincent Prévot, INSERM U1172, Lille). A vaginal smear was performed at the time of the experiment to check the stage of the estrous cycle. In all experiments, mice were fed a balanced standard diet (Chow Diet [CD]) (3.14 kcal/g; Safe R04; Safe Diet, Augy, France). Before sacrifice, plasma was collected by submandibular vein sampling in an EDTA tube (BD microtainer, K2E tubes) or Lithium Heparin tube (BD microtainer, Lithium Heparin tubes) and centrifuged (1500 g, 10min, 4°C) to recover the plasma stored at −80°C. Mice were killed by cervical dislocation. Then, organs were removed, weighed, if necessary, dissected and quickly frozen in liquid nitrogen. A liver sample was prepared for histological analysis. All *in vivo* experimental procedures were performed in accordance with the principles established by EU directive 2010/63/EU, the Institut National de la Santé et de la Recherche Médicale and approved by the local Ethical Committee of Animal Care (APAFIS 22956-2019112517144195v4).

### 2. *In vivo* experiments and nutritional settings

#### Circadian experiment with day restricted feeding (DRF)

Eleven-week old male and female *ERα^hep−/−^* and *ERα^hep+/+^* mice were fed *ad libitum* or forced to feed during the day (ZT0-ZT12) and fasted during the night (ZT12-ZT24) for 2 weeks (DRF). ZT0 and ZT12 correspond respectively to the time at which the light switches on and off in the animal facility. The mice were then euthanized every 4 hours for 24 hours from ZT0 to ZT24 (n=5 animals/sex/genotype/ZT).

#### Fasting/Feeding experiments

Eleven-week-old male and female *ERα^hep−/−^* and *ERα^hep+/+^* mice were fed *ad libitum* or fasted from ZT0, then euthanized at ZT16 (n= 6 animals/group) or fasted for 24 hours from ZT0 to ZT24 for ketonemia measurement (n=5/group).

### 3. Indirect calorimetry

Assessment of basal metabolism in 10-week old male and female *ERα^hep−/−^* and *ERα^hep+/+^* mice (n=4-7 animals/sex/genotype) was performed in metabolic cages using indirect calorimetry (Phenomaster System, TSE Systems) at the ANEXPLO/Genotoul Zootechnics Core Facilities in Toulouse (INSERM/CNRS/UPS/ENVT US006). Mice were acclimatized for 24 hours to single housing and feeding through the food hopper. Then, food and water intake, spontaneous locomotor activity and gas exchange were recorded for 24 hours (feeding *ad libitum*) or for 48 hours (24 hours of fasting followed by 24 hours of refeeding).

### 4. Biochemical analysis

Free fatty acids (FFA), total cholesterol (CHL), LDL cholesterol (LDLc), HDL cholesterol (HDLc), triglycerides and Alanine transaminase (ALT) were determined from plasma samples with a Pentra C400 Clinical Chemistry Analyzer (HORIBA Medical, France) at the ANEXPLO/Genotoul Zootechnics Core Facilities in Toulouse (INSERM/CNRS/UPS/ENVT US006). Glycemia and ketonemia (β-hydroxybutyrate) were measured on blood samples taken from the tail with Accu-Chek Performa (Roche Diabetes Care) and Freestyle Optium Neo (Abbott Laboratories), respectively. Plasma insulin was assayed using the rat/mouse insulin ELISA kit (EZRMR-13K, Merck Millipore) following the manufacturer’s instructions.

### 5. Western Blot

Using a Precellys tissue homogenizer, frozen tissues (50 mg) were homogenized in an ice-cold lysis buffer (50 mM TrisHCl pH 7.4, 150 mM NaCl, 1% NP-40, 0.25% sodium deoxycholate, 0.1% SDS, 2 mM EDTA) supplemented with a cocktail of protease and phosphatase inhibitors (Aprotinin, Leupeptin and sodium orthovanadate), then sonicated and centrifuged at 13,000 g for 10 minutes at 4°C. Total protein content was measured using BC Assay Protein Quantification Kit (Interchim) following the manufacturer’s instructions. After adding loading buffer, samples were heated at 95°C for 5 min and then stored at −20°C. Samples containing 40 µg total proteins were separated on polyacrylamide gel gradient (4-20%) (Invitrogen) and transferred to a nitrocellulose membrane (Transblot, Bio-Rad). After saturation, immunodetection was performed with an antibody directed against ERα (MC-20, SC-542, Santa Cruz, 1/1000). Revelation was performed using an HRP-conjugated secondary antibody (anti-rabbit HRP, 1/10 000, Cell Signaling) and visualized by ECL detection according to the manufacturer’s instructions (Amersham Biosciences/GE Healthcare), using ChemiDoc Touch (We-Met facility, I2MC, Toulouse). Finally, the densitometric analyses of the bands were performed using the Image Lab software (Bio-Rad).

### 6. Gene expression

RNAs were extracted from liver samples collected at the end of the experiments with 1mL of Trizol reagent (Invitrogen) in Lysing Matrix D (MP) bead tubes on a Precellys apparatus (Ozyme). RNAs were quantified using nanodrop (ThermoScientific Nanodrop1000, Les Ulis, France). Two micrograms of total RNA were reverse-transcribed using the High-Capacity cDNA Reverse Transcription Kit (Applied Biosystems) and cDNAs were diluted to the 20th and kept at −20°C. The primers used for cDNA amplification were designed using PrimerExpress software (Applied Biosystems), tested and validated before their use. The quantitative PCR reactions were carried out on 2μL of cDNA diluted to 20th with 5μL of SybrGreen (SsoFast EvaGreen Supermix, Bio-Rad) and 3μL of the pair of primers (300 nM final) (primer list, **Table S7**) on an ABI StepOnePlus Real Time PCR System thermocycler (ThermoFisher Scientific) (GeT-Santé facility, GenoToul, Génopole Toulouse Midi-Pyrénées). qPCR data were normalized to TATA-box-binding protein (TBP) mRNA levels and analyzed with LinRegPCR.v2015.3 to get the starting concentration (N0) which is calculated as follow: N0 = threshold/(Eff mean^Cq^) with Eff mean: mean PCR efficiency and Cq:quantification cycle. To assess rhythmicity and mean differences of gene expression in gene expression analysis we used the drylm function of the dryR (for Differential Rhythmicity Analysis in R package) built by Weger et al. 2021 (28): dryR: Differential RhythmicitY analysis; R package version 1.0.0., a model selection framework based on generalized linear models (GLMs).

### 7. Microarray gene expression studies

Gene expression profiles were obtained for five or six liver samples per group at the GeT-TRiX facility (GenoToul, Génopole Toulouse Midi-Pyrénées) using Agilent Sureprint G3 Mouse GE v2 microarrays (8×60K, design 074809) following the manufacturer’s instructions. For each sample, Cyanine-3 (Cy3) labelled cRNA was prepared from 200 ng of total RNA using the One-Color Quick Amp Labeling kit (Agilent) according to the manufacturer’s instructions, followed by Agencourt RNAClean XP (Agencourt Bioscience Corporation, Beverly, Massachusetts). Dye incorporation and cRNA yield were checked using a Dropsense™ 96 UV/VIS droplet reader (Trinean, Belgium). A total of 600 ng of Cy3-labelled cRNA was hybridized on the microarray slides following the manufacturer’s instructions. Immediately after washing, the slides were scanned on an Agilent G2505C Microarray Scanner using Agilent Scan Control A.8.5.1 software and the fluorescence signal extracted using Agilent Feature Extraction software v10.10.1.1 with default parameters. Microarray data and experimental details are available in NCBI’s Gene Expression Omnibus. Microarray data were analysed using Matrix application (http://matrix.toulouse.inra.fr/) (GeT-TRiX facility, GenoToul, Génopole Toulouse Midi-Pyrénées).

### 8. Correlation analysis

Correlation matrixes were created with gene expression levels. Genes were considered as correlated when expression correlation was found to be >0.8 with a p-value <0.05. These correlated genes were assembled in a correlation network. Transcriptional regulatory network was also built using Trrust (https://www.grnpedia.org/trrust/), a manually curated database of mouse transcriptional regulatory network providing information on the interactions between transcription factors (TFs) and their targets and insights into network hubs, motifs and hierarchical organization (Han et al., 2018). Networks of the 50 genes having the highest absolute correlation (Pearson-correlation) with a transcription factor of interest (red node) were displayed using the R function circlePlot54. Only the edges corresponding to significant correlations were represented (Bonferroni adjusted P value < 5%). Positive and negative correlations were represented by red and blue edges, respectively.

### 9. Reporter metabolite analysis

Reporter metabolite analyses were performed to investigate the metabolites affected by transcriptional changes in the absence of ERα using the PIANO package, which enriches the gene set analysis of genome-wide data by incorporating the directionality of gene expression and combining statistical hypotheses and methods in conjunction with a previously established genome-scale metabolic model of the liver, iHepatocyte2322 [42].

### 10. Statistical Analysis

Results are expressed as mean ± SEM. Analyses were performed using GraphPad Prism 7 software for Mac OS X (GraphPad Software, San Diego, CA, www.graphpad.com). Two-way ANOVA analyses were used to assess the interaction between the sex (Male *versus* Female) or the genotype (*ERα^hep−/−^ versus ERα^hep+/+^*) and the nutritional status (Fasted *versus* Fed). In the absence of significant interaction, statistical analyses regarding the respective influence of each parameter are provided. In case of significant interaction, Sidak’s post-tests were subsequently performed to compare the different groups two by two. Repeated measures ANOVA were used to compare changes over time between genotypes in metabolic cages. A p value of p<0.05 was considered statistically significant.

## Supplemental Figures Legend

**Supplemental Figure 1. Body weight, relative tissue weights and plasma lipid profile in mice fed *ad libitum* or submitted to DRF.** Twelve-weeks old male and synchronized female control (*ERα^hep+/+^*) mice were fed *ad libitum* or subjected to a daytime restricted feeding (DRF) during 2 weeks (*ERα^hep+/+^*) (n=5 mice/sex/genotype/ZT).**(A-D)** At the end of the experiment, body weight (A), relative liver (B), perigonadic (pg) (C) and sub-cutaneous (sc) (D) white adipose tissue (WAT) weights were measured. Data represent mean ± SEM, # sex effect, ### or §§§ p-value<0.001. **(E-H)** Quantification of plasma triglycerides (E), total cholesterol (F), HDL cholesterol (G) and LDL cholesterol (H) levels determined from plasma samples collected at the end of the experiment. Data were analyzed with drylm function from the dryR package. Groups sharing the same fitting line shape have the same rhythmic parameters. A flat line signifies no rhythm detected.

**Supplemental Figure 2. Hepatocyte ERα deletion has no impact on body weight, relative tissue weights and plasma lipid profile in male and female mice fed *ad libitum* or under DRF.** Twelve-weeks old male and synchronized female hepatocyte-specific knockout mice (*ERα^hep−/−^*) and their control littermates (*ERα^hep+/+^*) were fed *ad libitum* or subjected to a day-restricted feeding (DRF, ZT0-ZT12) during 2 weeks (n=5 mice/sex/genotype/ZT). **(A-C)** At the end of the experiment, body weight (A), relative liver (B) and perigonadic white adipose tissue (pgWAT) (C) weights were measured. Data represent mean ± SEM. * genotype effect, ** or §§ p-value<0.01. **(D-G)** Quantification of plasma triglycerides (D), total cholesterol (E), HDLc (F) and LDLc (G) levels determined from plasma samples collected at the end of the experiment. Data were analyzed with drylm function from the dryR package. Groups sharing the same fitting line shape have the same rhythmic parameters. A flat line signifies no rhythm detected.

**Supplemental Figure 3. Validation of nutritional status in mice fed *ad libitum* or fasted for 16 hours**. Twelve week-old male and synchronized female ERα hepatocyte-specific deleted mice (*ERα^hep−/−^*) and their control littermates (*ERα^hep+/+^*) were fed *ad libitum* or fasted for 16h and then killed at ZT16 (n=6/group). **(A-C)** Quantification of circulating glucose (A), insulin (B) and FFA (C) plasma levels at the end of the experiment. **(D-E)** Relative liver gene expression of *G6pc* (D) and *Srebp1c* (E) determined by qRT-PCR. TBP was used as housekeeping gene. Data represent mean ± SEM. Statistical significance was calculated using two-way ANOVA with Sidak’s multiple comparisons. * genotype effect, § nutritional effect. * or § p-value < 0.05; ** or §§ p-value < 0.01; *** or §§§ p-value < 0.001.

**Supplemental Figure 4. Correlation network of *ERα* is sex-dimorphic in Human and mouse. (A-B)** Correlation network of *ESR1* in Human (A) and in mouse (B) established from GSE159088 and GSE159090, respectively. The edges corresponding to significant correlations are presented (Bonferroni-adjusted p<0.05). The network circle plot based on the top 30 men/male biased genes on the right (blue nodes) and on the top 30 women/female biased genes on the left (magenta nodes). Correlations with these selected genes in men/male is presented on the left while the corresponding circle plot in women/female is on the right. Red and blue were used for positive and negative correlations, respectively.

**Supplemental Figure 5. Principal Component Analysis established from liver transcriptome in fed and fasted *ERα^hep+/+^* and *ERα^hep−/−^* females and males.** Twelve week-old male and synchronized female ERα hepatocyte-specific deleted mice (*ERα^hep−/−^*) and their littermate controls (*ERα^hep+/+^*) were fed *ad libitum* or fasted for 16h and then killed at ZT16 (n=6/group). Liver transcriptome was determined by microarrays. **(A-B)** Principal component analysis from microarrays data in fed and fasted female (A) and male (B) *ERα^hep+/+^* and *ERα^hep−/−^* mice.

**Supplemental Figure 6. Volcano plot established from transcriptomic data from fasted and fed female *ERα^hep+/+^* mice (n=6/group).**

**Supplemental Figure 7. Hepatocyte ERα-dependent sex-biased genes in liver of female mice.** Twelve-weeks old synchronized female hepatocyte-specific knockout mice (*ERα^hep−/−^*) and their littermate controls (*ERα^hep+/+^*) were fed *ad libitum* or fasted for 16h and then killed at ZT16 (n=6/group). Liver transcriptome was analyzed by microarrays. **(A)** Hepatocyte ERα-dependent SBG in female (p-value<0.05, |FC|>2) was established from liver transcriptomic data. **(B)** Gene Ontology analysis performed on the 237 hepatocyte ERα-dependent sex-biased genes was performed and transcription factors involved established from TTRUST database.

**Supplemental Figure 7. In both sexes, hepatocyte ERα has no influence on metabolic parameters.** Metabolic parameters were registered in metabolic cages in male and synchronized female hepatocyte-specific knockout mice (*ERα^hep−/−^*) and their littermate controls (*ERα^hep+/+^*) fed *ad libitum* during 24 hours. **(A-B)** Respiratory energy rate evolution during the 24 hours recorded in male (A) and female (B). Data represent mean ± SEM for each ZT; Repeated measures ANOVA were used to compare changes over time between the genotypes. **(C-F)** Mean values of RER (C), energy expenditure (D), activity (E) and food intake (F) are shown according to the sex and the day (light) and night (dark) periods. Data represent mean ± SEM; statistical significance was calculated using two-way ANOVA with Sidak’s multiple comparisons.

## References

1. Roy AK, Chatterjee B. Sexual dimorphism in the liver. Annu Rev Physiol. 1983;45:37–50.

2. Yang X, Schadt EE, Wang S, Wang H, Arnold AP, Ingram-Drake L, et al. Tissue-specific expression and regulation of sexually dimorphic genes in mice. Genome Res. août 2006;16(8):995–1004.

3. Goossens GH, Jocken JWE, Blaak EE. Sexual dimorphism in cardiometabolic health: the role of adipose tissue, muscle and liver. Nat Rev Endocrinol. janv 2021;17(1):47–66.

4. Waxman DJ, Holloway MG. Sex differences in the expression of hepatic drug metabolizing enzymes. Mol Pharmacol. août 2009;76(2):215–28.

5. Maggi A, Della Torre S. Sex, metabolism and health. Mol Metab. sept 2018;15:3–7.

6. Goldfarb CN, Karri K, Pyatkov M, Waxman DJ. Interplay Between GH-regulated, Sex-biased Liver Transcriptome and Hepatic Zonation Revealed by Single-Nucleus RNA Sequencing. Endocrinology. 1 juill 2022;163(7):bqac059.

7. Lefebvre P, Staels B. Hepatic sexual dimorphism -implications for non-alcoholic fatty liver disease. Nat Rev Endocrinol. nov 2021;17(11):662–70.

8. van Nas A, Guhathakurta D, Wang SS, Yehya N, Horvath S, Zhang B, et al. Elucidating the role of gonadal hormones in sexually dimorphic gene coexpression networks. Endocrinology. mars 2009;150(3):1235–49.

9. Della Torre S. Non-alcoholic Fatty Liver Disease as a Canonical Example of Metabolic Inflammatory-Based Liver Disease Showing a Sex-Specific Prevalence: Relevance of Estrogen Signaling. Front Endocrinol (Lausanne). 2020;11:572490.

10. Blencowe M, Chen X, Zhao Y, Itoh Y, McQuillen CN, Han Y, et al. Relative contributions of sex hormones, sex chromosomes, and gonads to sex differences in tissue gene regulation. Genome Res. mai 2022;32(5):807–24.

11. Lonardo A, Nascimbeni F, Ballestri S, Fairweather D, Win S, Than TA, et al. Sex Differences in Nonalcoholic Fatty Liver Disease: State of the Art and Identification of Research Gaps. Hepatology. oct 2019;70(4):1457–69.

12. Tramunt B, Smati S, Grandgeorge N, Lenfant F, Arnal JF, Montagner A, et al. Sex differences in metabolic regulation and diabetes susceptibility. Diabetologia. mars 2020;63(3):453–61.

13. Mauvais-Jarvis F, Clegg DJ, Hevener AL. The role of estrogens in control of energy balance and glucose homeostasis. Endocr Rev. juin 2013;34(3):309–38.

14. Morselli E, Santos RS, Criollo A, Nelson MD, Palmer BF, Clegg DJ. The effects of oestrogens and their receptors on cardiometabolic health. Nat Rev Endocrinol. juin 2017;13(6):352–64.

15. Zhu L, Brown WC, Cai Q, Krust A, Chambon P, McGuinness OP, et al. Estrogen treatment after ovariectomy protects against fatty liver and may improve pathway-selective insulin resistance. Diabetes. févr 2013;62(2):424–34.

16. Lemieux C, Phaneuf D, Labrie F, Giguère V, Richard D, Deshaies Y. Estrogen receptor alpha-mediated adiposity-lowering and hypocholesterolemic actions of the selective estrogen receptor modulator acolbifene. Int J Obes (Lond). oct 2005;29(10):1236–44.

17. Handgraaf S, Riant E, Fabre A, Waget A, Burcelin R, Lière P, et al. Prevention of obesity and insulin resistance by estrogens requires ERα activation function-2 (ERαAF-2), whereas ERαAF-1 is dispensable. Diabetes. déc 2013;62(12):4098–108.

18. Della Torre S. Beyond the X Factor: Relevance of Sex Hormones in NAFLD Pathophysiology. Cells. 21 sept 2021;10(9):2502.

19. Guillaume M, Riant E, Fabre A, Raymond-Letron I, Buscato M, Davezac M, et al. Selective Liver Estrogen Receptor α Modulation Prevents Steatosis, Diabetes, and Obesity Through the Anorectic Growth Differentiation Factor 15 Hepatokine in Mice. Hepatol Commun. juill 2019;3(7):908–24.

20. Villa A, Della Torre S, Stell A, Cook J, Brown M, Maggi A. Tetradian oscillation of estrogen receptor α is necessary to prevent liver lipid deposition. Proc Natl Acad Sci U S A. 17 juill 2012;109(29):11806–11.

21. Della Torre S, Rando G, Meda C, Stell A, Chambon P, Krust A, et al. Amino acid-dependent activation of liver estrogen receptor alpha integrates metabolic and reproductive functions via IGF-1. Cell Metab. 2 févr 2011;13(2):205–14.

22. Della Torre S, Mitro N, Fontana R, Gomaraschi M, Favari E, Recordati C, et al. An Essential Role for Liver ERα in Coupling Hepatic Metabolism to the Reproductive Cycle. Cell Rep. 12 avr 2016;15(2):360–71.

23. Asher G, Sassone-Corsi P. Time for food: the intimate interplay between nutrition, metabolism, and the circadian clock. Cell. 26 mars 2015;161(1):84–92.

24. Mermet J, Yeung J, Naef F. Systems Chronobiology: Global Analysis of Gene Regulation in a 24-Hour Periodic World. Cold Spring Harb Perspect Biol. 1 mars 2017;9(3):a028720.

25. Reinke H, Asher G. Crosstalk between metabolism and circadian clocks. Nat Rev Mol Cell Biol. avr 2019;20(4):227–41.

26. Bolshette N, Ibrahim H, Reinke H, Asher G. Circadian regulation of liver function: from molecular mechanisms to disease pathophysiology. Nat Rev Gastroenterol Hepatol. nov 2023;20(11):695–707.

27. Damiola F, Le Minh N, Preitner N, Kornmann B, Fleury-Olela F, Schibler U. Restricted feeding uncouples circadian oscillators in peripheral tissues from the central pacemaker in the suprachiasmatic nucleus. Genes Dev. 1 déc 2000;14(23):2950–61.

28. Weger BD, Gobet C, David FPA, Atger F, Martin E, Phillips NE, et al. Systematic analysis of differential rhythmic liver gene expression mediated by the circadian clock and feeding rhythms. Proc Natl Acad Sci U S A. 19 janv 2021;118(3):e2015803118.

29. Blattner MS, Mahoney MM. Photic phase-response curve in 2 strains of mice with impaired responsiveness to estrogens. J Biol Rhythms. août 2013;28(4):291–300.

30. Alvord VM, Kantra EJ, Pendergast JS. Estrogens and the circadian system. Semin Cell Dev Biol. juin 2022;126:56–65.

31. Aizarani N, Saviano A, Sagar null, Mailly L, Durand S, Herman JS, et al. A human liver cell atlas reveals heterogeneity and epithelial progenitors. Nature. août 2019;572(7768):199–204.

32. Montagner A, Polizzi A, Fouché E, Ducheix S, Lippi Y, Lasserre F, et al. Liver PPARα is crucial for whole-body fatty acid homeostasis and is protective against NAFLD. Gut. juill 2016;65(7):1202–14.

33. Bideyan L, Nagari R, Tontonoz P. Hepatic transcriptional responses to fasting and feeding. Genes Dev. 1 mai 2021;35(9-10):635–57.

34. Smati S, Polizzi A, Fougerat A, Ellero-Simatos S, Blum Y, Lippi Y, et al. Integrative study of diet-induced mouse models of NAFLD identifies PPARα as a sexually dimorphic drug target. Gut. avr 2022;71(4):807–21.

35. Mardinoglu A, Shoaie S, Bergentall M, Ghaffari P, Zhang C, Larsson E, et al. The gut microbiota modulates host amino acid and glutathione metabolism in mice. Mol Syst Biol. 16 oct 2015;11(10):834.

36. Eagon PK, DiLeo A, Polimeno L, Francavilla A, Van Thiel DH, Guglielmi F, et al. Circadian rhythm of hepatic cytosolic and nuclear estrogen receptors. Chronobiol Int. 1986;3(4):207–11.

37. Kornmann B, Schaad O, Reinke H, Saini C, Schibler U. Regulation of circadian gene expression in liver by systemic signals and hepatocyte oscillators. Cold Spring Harb Symp Quant Biol. 2007;72:319–30.

38. Vollmers C, Gill S, DiTacchio L, Pulivarthy SR, Le HD, Panda S. Time of feeding and the intrinsic circadian clock drive rhythms in hepatic gene expression. Proc Natl Acad Sci U S A. 15 déc 2009;106(50):21453–8.

39. Bingel AS, Schwartz NB. Timing of LH release and ovulation in the cyclic mouse. J Reprod Fertil. juill 1969;19(2):223–9.

40. Legan SJ, Coon GA, Karsch FJ. Role of estrogen as initiator of daily LH surges in the ovariectomized rat. Endocrinology. janv 1975;96(1):50–6.

41. Gery S, Virk RK, Chumakov K, Yu A, Koeffler HP. The clock gene Per2 links the circadian system to the estrogen receptor. Oncogene. 13 déc 2007;26(57):7916–20.

42. Xiao L, Chang AK, Zang MX, Bi H, Li S, Wang M, et al. Induction of the CLOCK gene by E2-ERα signaling promotes the proliferation of breast cancer cells. PLoS One. 2014;9(5):e95878.

43. Palmisano BT, Stafford JM, Pendergast JS. High-Fat Feeding Does Not Disrupt Daily Rhythms in Female Mice because of Protection by Ovarian Hormones. Front Endocrinol (Lausanne). 2017;8:44.

44. Omotola O, Legan S, Slade E, Adekunle A, Pendergast JS. Estradiol regulates daily rhythms underlying diet-induced obesity in female mice. Am J Physiol Endocrinol Metab. 1 déc 2019;317(6):E1172–81.

45. Racanelli V, Rehermann B. The liver as an immunological organ. Hepatology. févr 2006;43(2 Suppl 1):S54–62.

46. Heymann F, Tacke F. Immunology in the liver--from homeostasis to disease. Nat Rev Gastroenterol Hepatol. févr 2016;13(2):88–110.

47. Hotamisligil GS. Inflammation and metabolic disorders. Nature. 14 déc 2006;444(7121):860–7.

48. Robinson MW, Harmon C, O’Farrelly C. Liver immunology and its role in inflammation and homeostasis. Cell Mol Immunol. mai 2016;13(3):267–76.

49. Frasor J, El-Shennawy L, Stender JD, Kastrati I. NFκB affects estrogen receptor expression and activity in breast cancer through multiple mechanisms. Mol Cell Endocrinol. 15 déc 2015;418 Pt 3(0 3):235–9.

50. Björnström L, Sjöberg M. Mechanisms of estrogen receptor signaling: convergence of genomic and nongenomic actions on target genes. Mol Endocrinol. avr 2005;19(4):833–42.

51. Goldstein I, Hager GL. Transcriptional and Chromatin Regulation during Fasting -The Genomic Era. Trends Endocrinol Metab. déc 2015;26(12):699–710.

52. Strable MS, Ntambi JM. Genetic control of de novo lipogenesis: role in diet-induced obesity. Crit Rev Biochem Mol Biol. juin 2010;45(3):199–214.

53. Han S iee, Komatsu Y, Murayama A, Steffensen KR, Nakagawa Y, Nakajima Y, et al. Estrogen receptor ligands ameliorate fatty liver through a nonclassical estrogen receptor/Liver X receptor pathway in mice. Hepatology. mai 2014;59(5):1791–802.

54. Leuenberger N, Pradervand S, Wahli W. Sumoylated PPARalpha mediates sex-specific gene repression and protects the liver from estrogen-induced toxicity in mice. J Clin Invest. oct 2009;119(10):3138–48.

55. Huss JM, Torra IP, Staels B, Giguère V, Kelly DP. Estrogen-related receptor alpha directs peroxisome proliferator-activated receptor alpha signaling in the transcriptional control of energy metabolism in cardiac and skeletal muscle. Mol Cell Biol. oct 2004;24(20):9079–91.

56. Bonofiglio D, Gabriele S, Aquila S, Catalano S, Gentile M, Middea E, et al. Estrogen receptor alpha binds to peroxisome proliferator-activated receptor response element and negatively interferes with peroxisome proliferator-activated receptor gamma signaling in breast cancer cells. Clin Cancer Res. 1 sept 2005;11(17):6139–47.

57. Tiano JP, Mauvais-Jarvis F. Molecular mechanisms of estrogen receptors’ suppression of lipogenesis in pancreatic β-cells. Endocrinology. juill 2012;153(7):2997–3005.

58. Pedram A, Razandi M, Blumberg B, Levin ER. Membrane and nuclear estrogen receptor α collaborate to suppress adipogenesis but not triglyceride content. FASEB J. janv 2016;30(1):230–40.

59. Lu Y, Tian N, Hu L, Meng J, Feng M, Zhu Y, et al. ERα down-regulates carbohydrate responsive element binding protein and decreases aerobic glycolysis in liver cancer cells. J Cell Mol Med. avr 2021;25(7):3427–36.

60. Palierne G, Fabre A, Solinhac R, Le Péron C, Avner S, Lenfant F, et al. Changes in Gene Expression and Estrogen Receptor Cistrome in Mouse Liver Upon Acute E2 Treatment. Mol Endocrinol. juill 2016;30(7):709–32.

