## Supplemental Figures for "Hepatocyte ERα orchestrates sex-specific liver adaptation to fasting and feeding states"

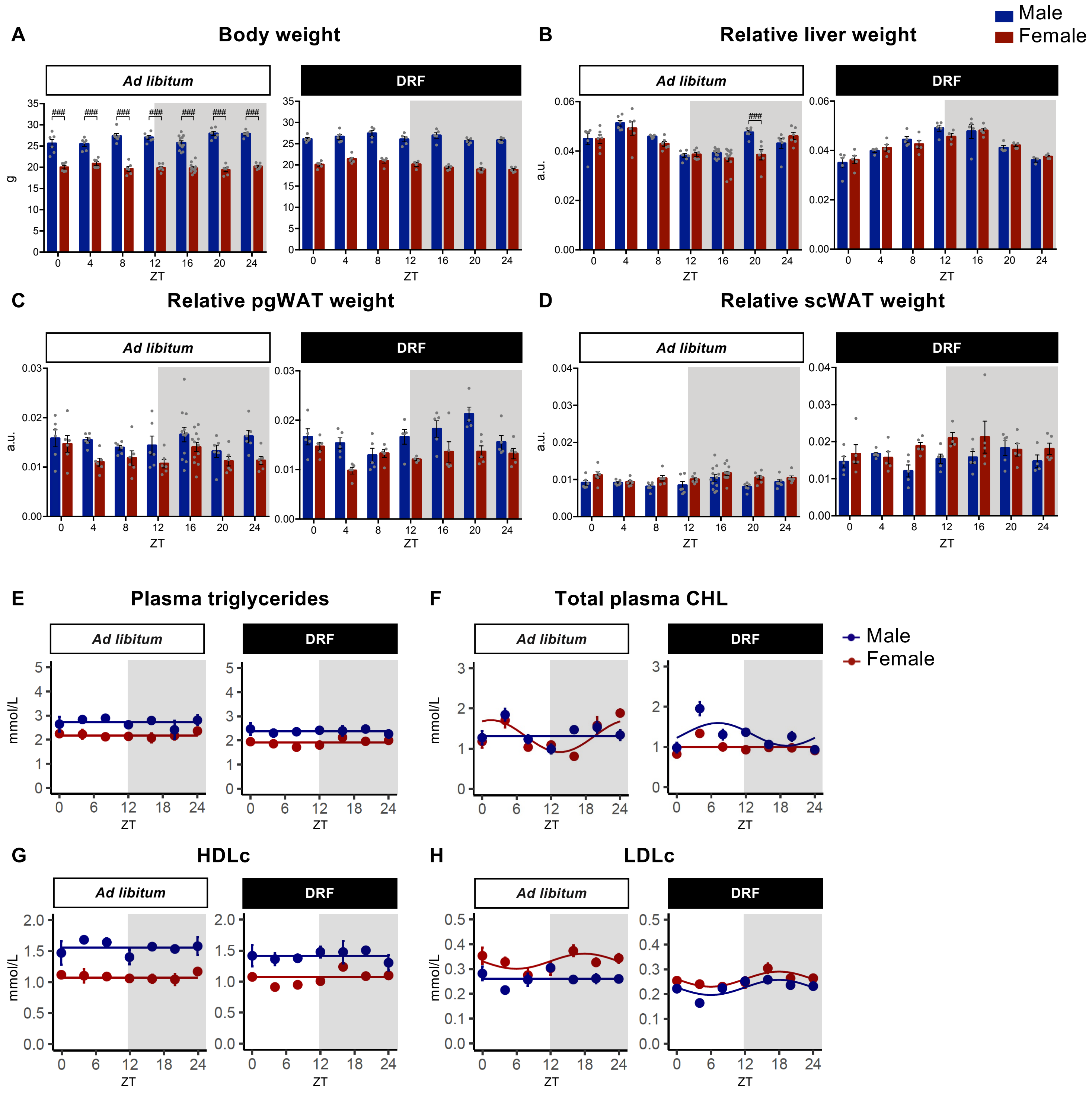
A
Body weight
B
Relative liver weight
Male
Female
Ad libitum
DRF
###
g
a.u.
ZT
C
Relative pgWAT weight
D
Relative scWAT weight
E
Plasma triglycerides
F
Total plasma CHL
G
HDLc
H
LDLc
mmol/L

Fig.S1

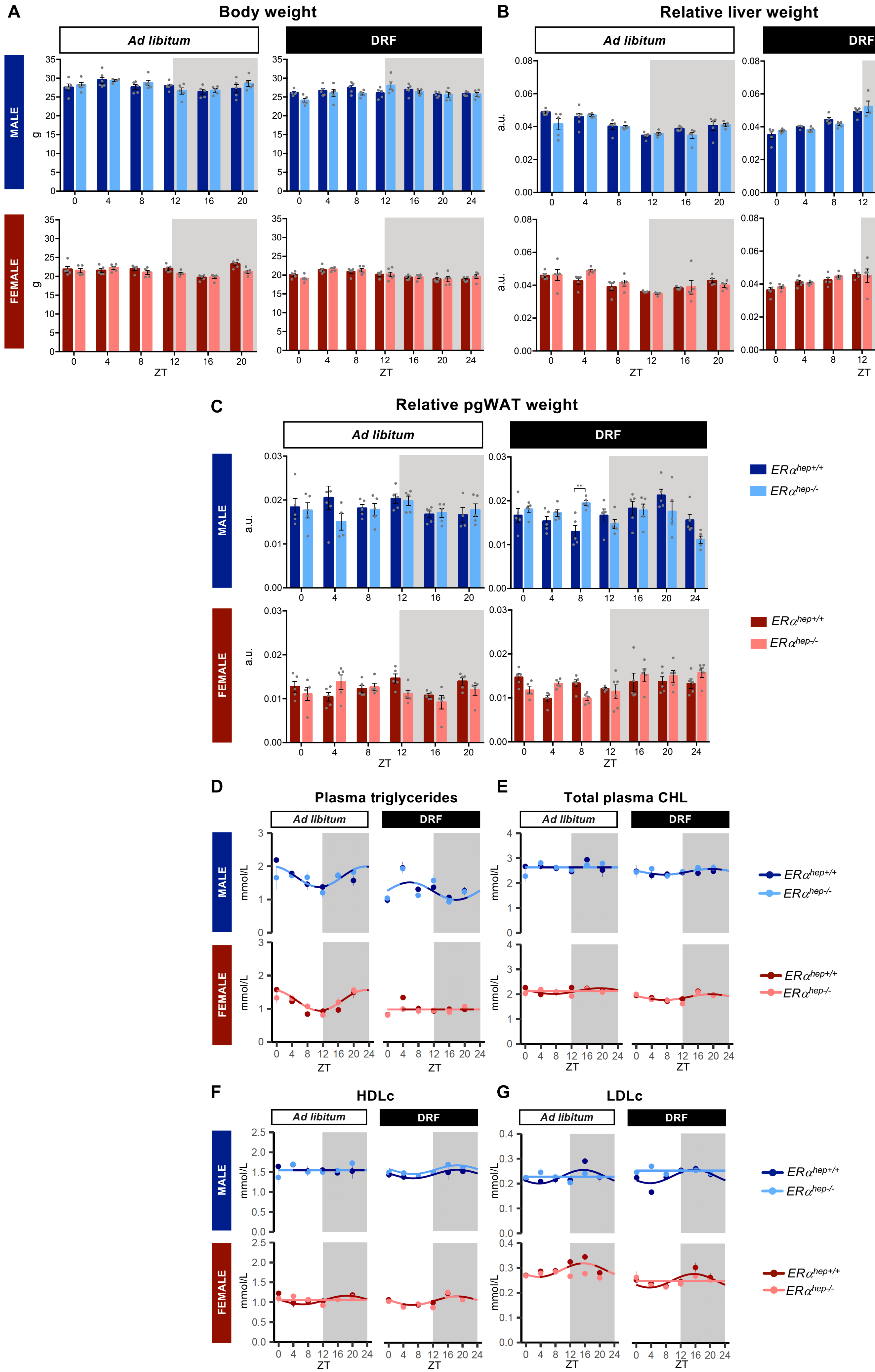
A
Body weight
Ad libitum
DRF
MALE
FEMALE
g
ZT
B
Relative liver weight
a.u.
C
Relative pgWAT weight
**
D
Plasma triglycerides
mmol/L
E
Total plasma CHL
F
HDLc
G
LDLc
ERα hep+/+
ERα hep-/-

Fig.S2

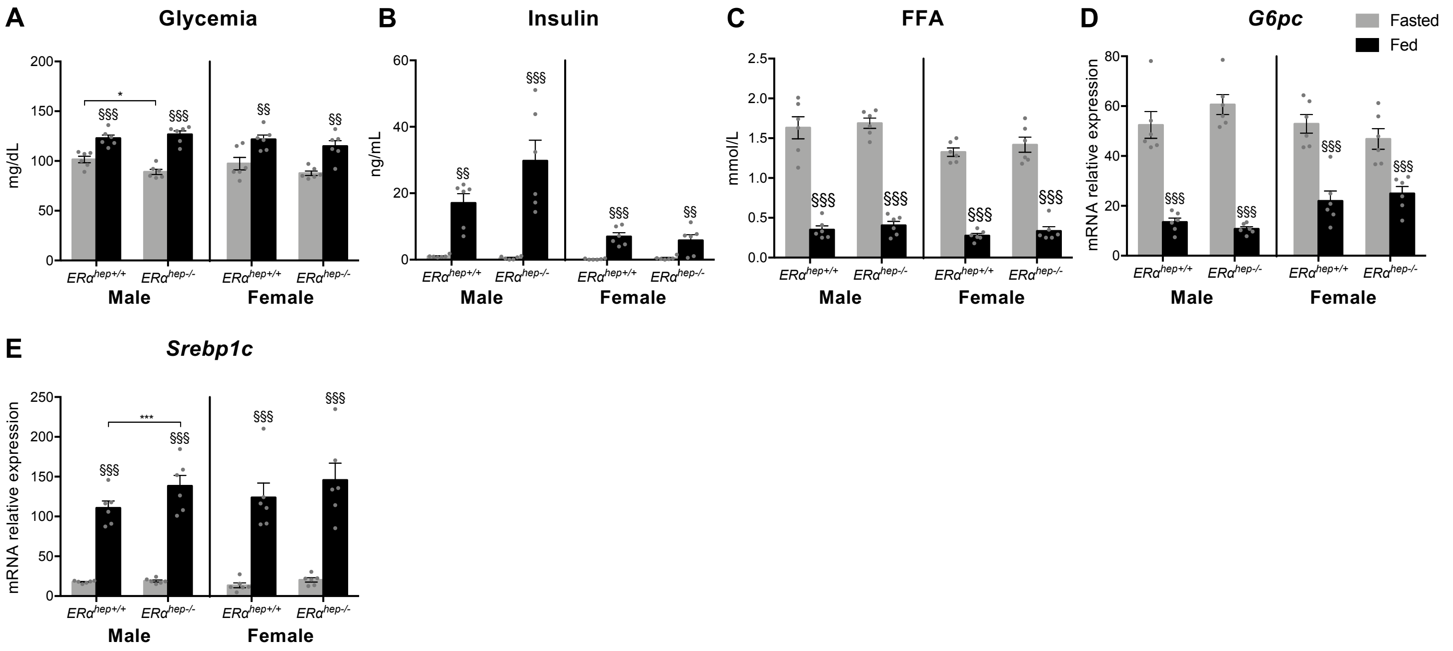
A
Glycemia
mg/dL
B
Insulin
ng/mL
C
FFA
mmol/L
D
G6pc
mRNA relative expression
Fasted
Fed
E
Srebp1c
mRNA relative expression
ERα hep+/+
ERα hep-/-
Male
Female

**Fig.S3**

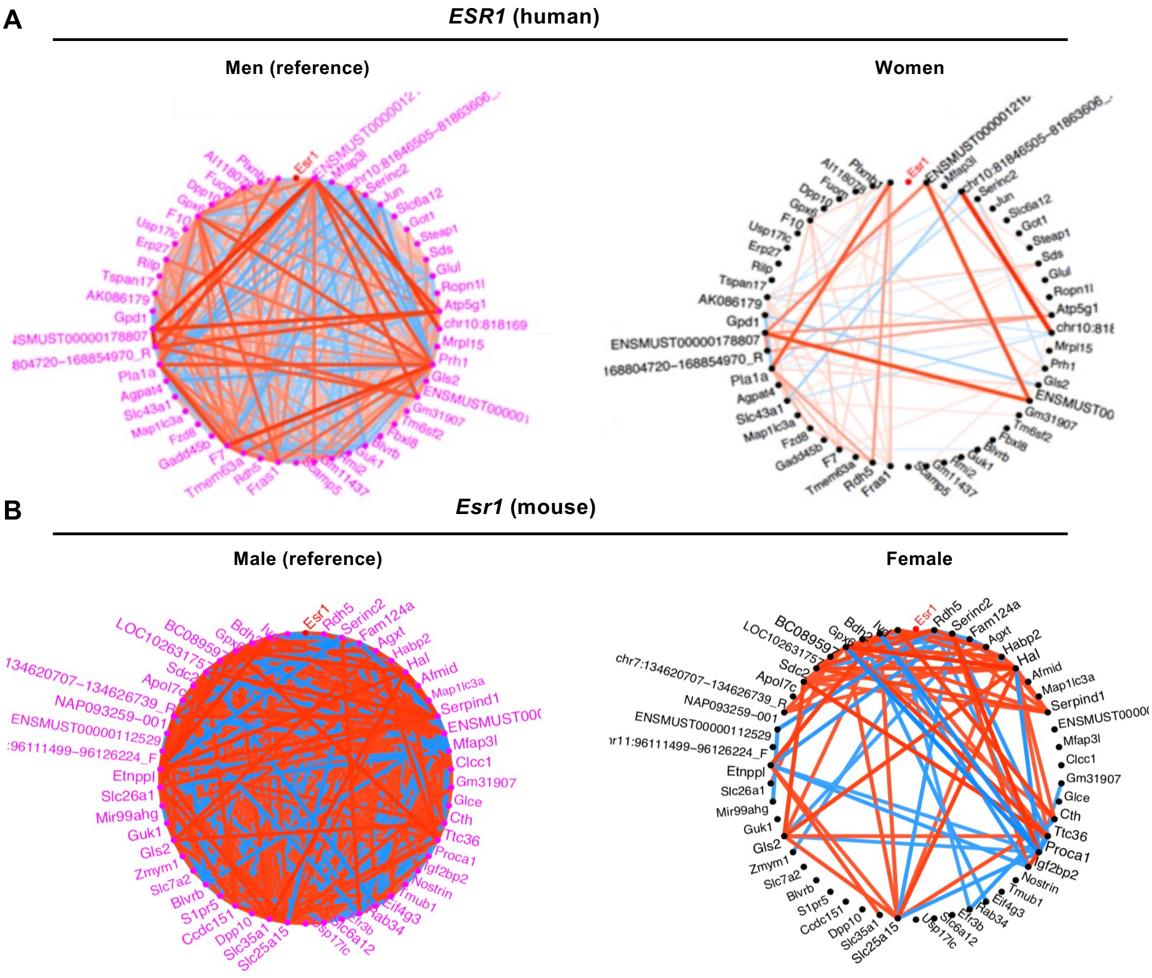


Genes the most correlated to *Esr1*

Top 30 genes in Men/Male

Top 30 genes in Women/Female

positive correlation

negative correlation

**Fig.S4**

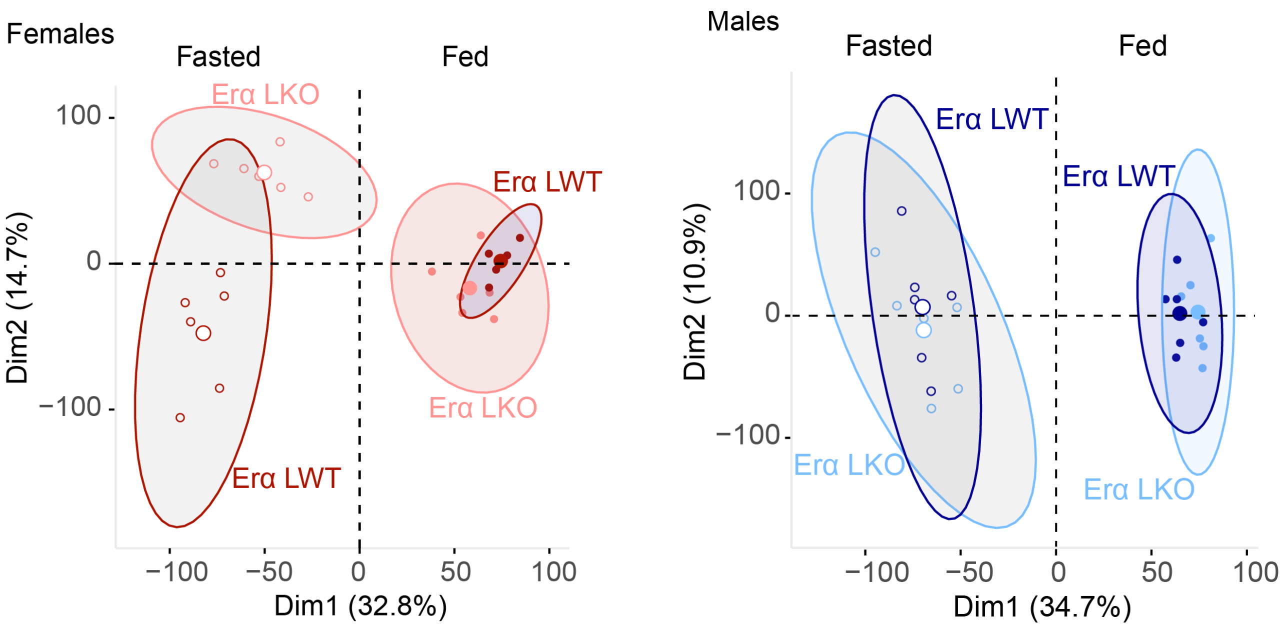
Females
Fasted
Fed
Erα LKO
Erα LWT
Erα LKO
Erα LWT
100
0
−100
Dim2 (14.7%)
−100
−50
0
50
100
Dim1 (32.8%)
Males
Fasted
Fed
Erα LWT
Erα LWT
Erα LKO
Erα LKO
100
0
−100
Dim2 (10.9%)
−100
−50
0
50
100
Dim1 (34.7%)

**Fig.S5**

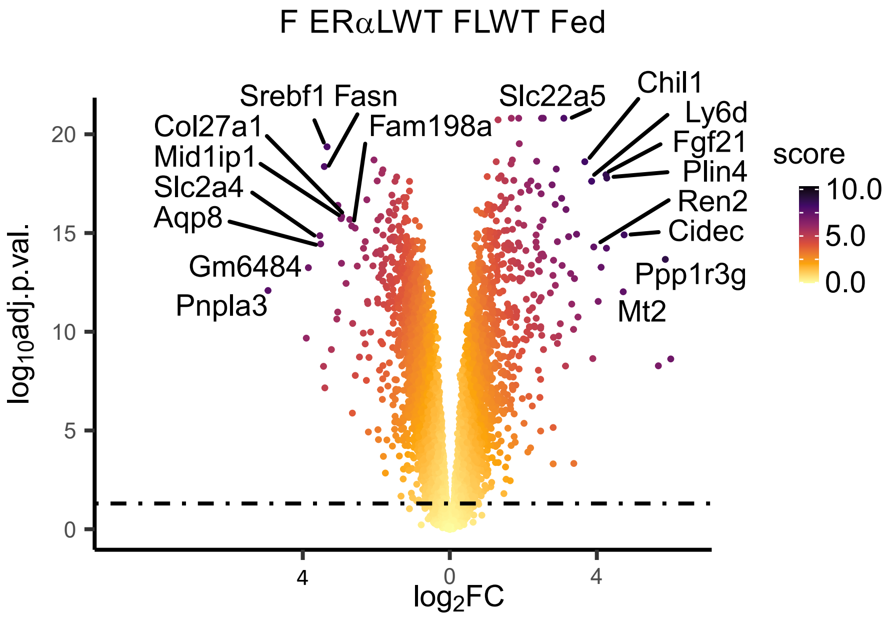

score = sqrt(abs(-log10adj.p.val*logFC))

**Fig.S6**

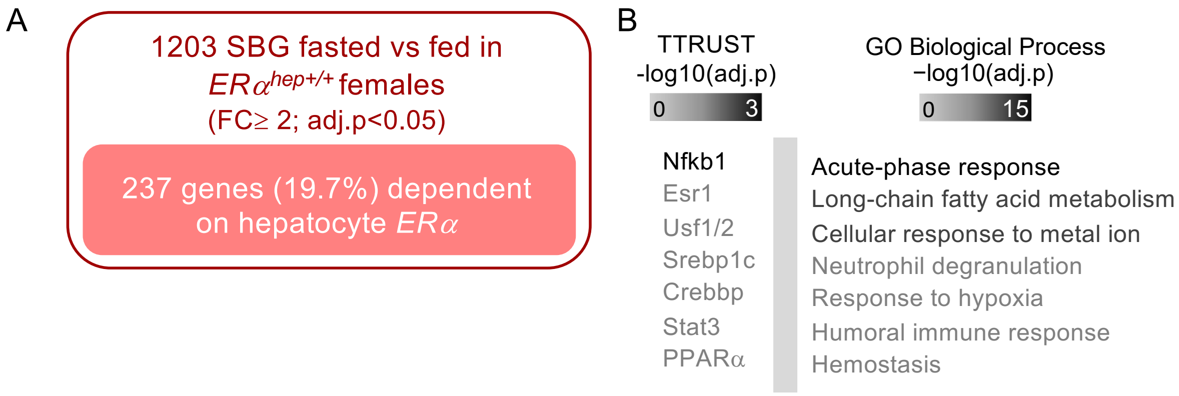
A
1203 SBG fasted vs fed in
ERα hep+/+ females
(FC≥ 2; adj.p<0.05)
237 genes (19.7%) dependent on hepatocyte ERα
B
TTRUST
-log10(adj.p)
0
3
GO Biological Process
−log10(adj.p)
0
15
Nfkb1
Esr1
Usf1/2
Srebp1c
Crebbp
Stat3
PPARα
Acute-phase response
Long-chain fatty acid metabolism
Cellular response to metal ion
Neutrophil degranulation
Response to hypoxia
Humoral immune response
Hemostasis

**Fig.S7**

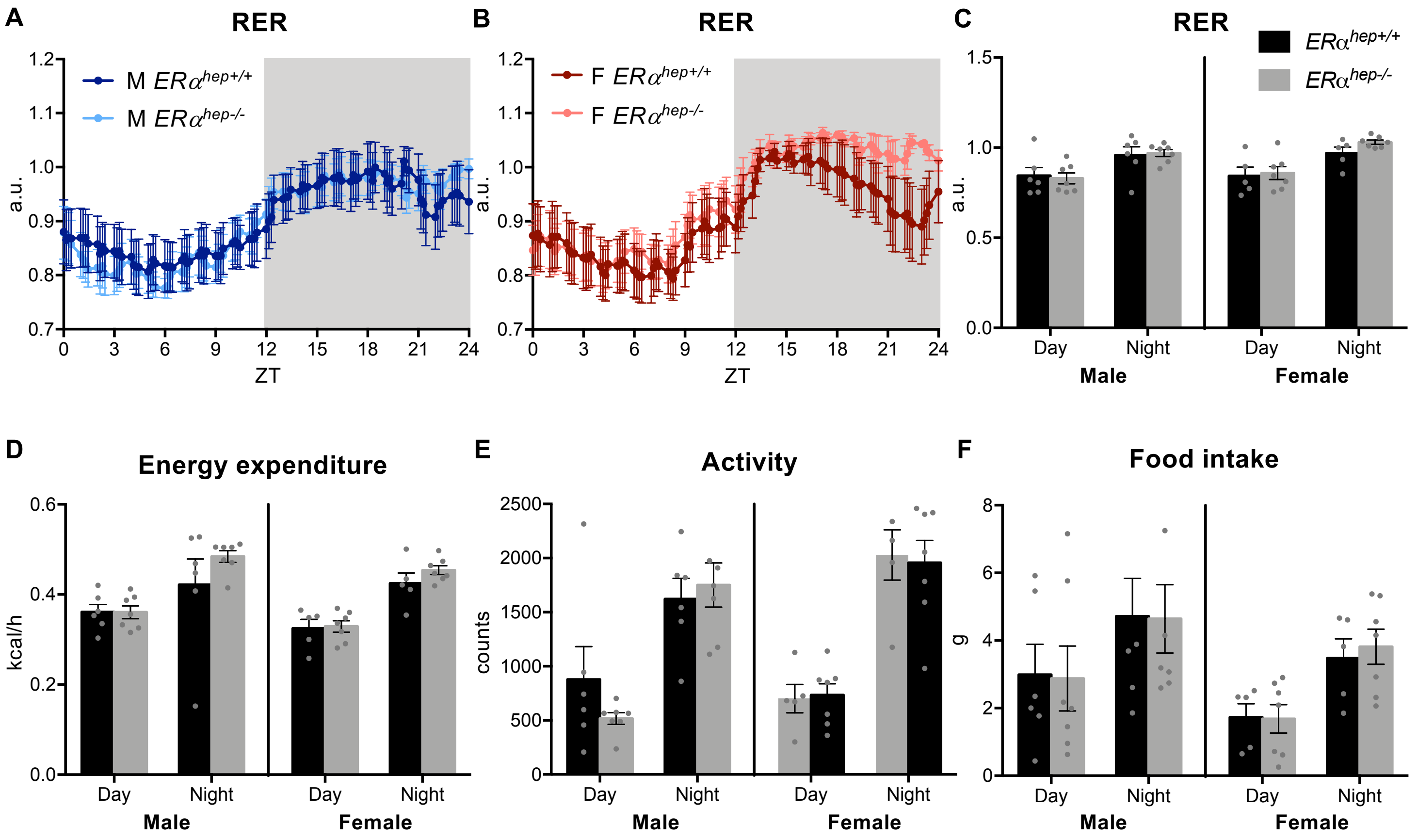

A
RER
M ERα hep+/+
M ERα hep-/-
a.u.
ZT
B
RER
F ERα hep+/+
F ERα hep-/-
C
RER
ERα hep+/+
ERα hep-/-
Day
Night
Male
Female
D
Energy expenditure
kcal/h
E
Activity
counts
F
Food intake
g


**Fig. S8**
